# Multimodal memory T cell profiling identifies a reduction in a polyfunctional Th17 state associated with tuberculosis progression

**DOI:** 10.1101/2020.04.23.057828

**Authors:** Aparna Nathan, Jessica I. Beynor, Yuriy Baglaenko, Sara Suliman, Kazuyoshi Ishigaki, Samira Asgari, Chuan-Chin Huang, Yang Luo, Zibiao Zhang, Kattya Lopez Tamara, Judith Jimenez, Roger I. Calderón, Leonid Lecca, Ildiko van Rhijn, D. Branch Moody, Megan B. Murray, Soumya Raychaudhuri

## Abstract

*Mycobacterium tuberculosis* (*M*.*tb*) results in 10 million active tuberculosis (TB) cases and 1.5 million deaths each year^1^, making it the world’s leading infectious cause of death^2^. Infection leads to either an asymptomatic latent state or TB disease. Memory T cells have been implicated in TB disease progression, but the specific cell states involved have not yet been delineated because of the limited scope of traditional profiling strategies. Furthermore, immune activation during infection confounds underlying differences in T cell state distributions that influence risk of progression. Here, we used a multimodal single-cell approach to integrate measurements of transcripts and 30 functionally relevant surface proteins to comprehensively define the memory T cell landscape at steady state (i.e., outside of active infection). We profiled 500,000 memory T cells from 259 Peruvians > 4.7 years after they had either latent *M*.*tb* infection or active disease and defined 31 distinct memory T cell states, including a CD4+CD26+CD161+CCR6+ effector memory state that was significantly reduced in patients who had developed active TB (OR = 0.80, 95% CI: 0.73–0.87, p = 1.21 × 10^−6^). This state was also polyfunctional; in *ex vivo* stimulation, it was enriched for IL-17 and IL-22 production, consistent with a Th17-skewed phenotype, but also had more capacity to produce IFNγ than other CD161+CCR6+ Th17 cells. Additionally, in progressors, IL-17 and IL-22 production in this cell state was significantly lower than in non-progressors. Reduced abundance and function of this state may be an important factor in failure to control *M*.*tb* infection.

## Main text

Interindividual immune differences may underlie host variation in response to pathogens, such as *M*.*tb*. Only 5-15% of individuals who are infected with *M*.*tb* develop TB disease during their lifetime^2^. Disease progression is influenced by host immune and genetic factors that implicate T cells, which are major contributors to defense against intracellular pathogens^3-11^. These markedly different outcomes of infection raise the question of whether steady-state differences in T cell composition underlie divergent host response to *M*.*tb*.

Studies examining immunophenotypes in active TB have identified numerous memory T cell changes, including in the CD4+^12^, activated^13^, exhausted^14-16^, Th1^17,18^, and IL-17+ compartments^19-23^. However, these studies typically profiled patients during ongoing infection and focused on antigen-specific T cells, rather than broad, intrinsic differences in memory T cell composition outside of acute infection or active disease. Moreover, it is challenging to acquire an adequate sample size, account for confounders influencing T cell composition^24^, and overcome limitations of surface marker or bulk RNA-seq-based technologies that only capture certain cell state changes.

Here, we profiled total memory T cells at single-cell resolution from patients more than four years after TB disease in order to identify broad steady-state differences in progressors with minimal interference from acute immune response. We re-recruited 259 individuals from a larger epidemiological study (n = 14,044) in Lima, Peru that identified patients with active TB disease and followed their *M*.*tb*-infected household contacts for one year to monitor progression to active disease (**Fig. 1**)^25^. Participants who were diagnosed with microbiologically confirmed TB were classified as cases; household contacts who were tuberculin skin test (TST)-positive and had not developed TB disease by time of re-recruitment (4.72–6.60 years [median: 5.7] after initial recruitment) were classified as controls. During this time, cases were treated for active disease, which has an estimated cure rate of at least 95%, so they were expected to return to an immune steady state^12,26^. The larger epidemiological cohort has been comprehensively characterized with a variety of socioeconomic and demographic traits. In our subset, TB progression was associated with host factors, such as age, height, weight, sex, and body mass index (BMI), consistent with the larger cohort (**Supplementary Table 1** and **2**).

**Fig. 1.**
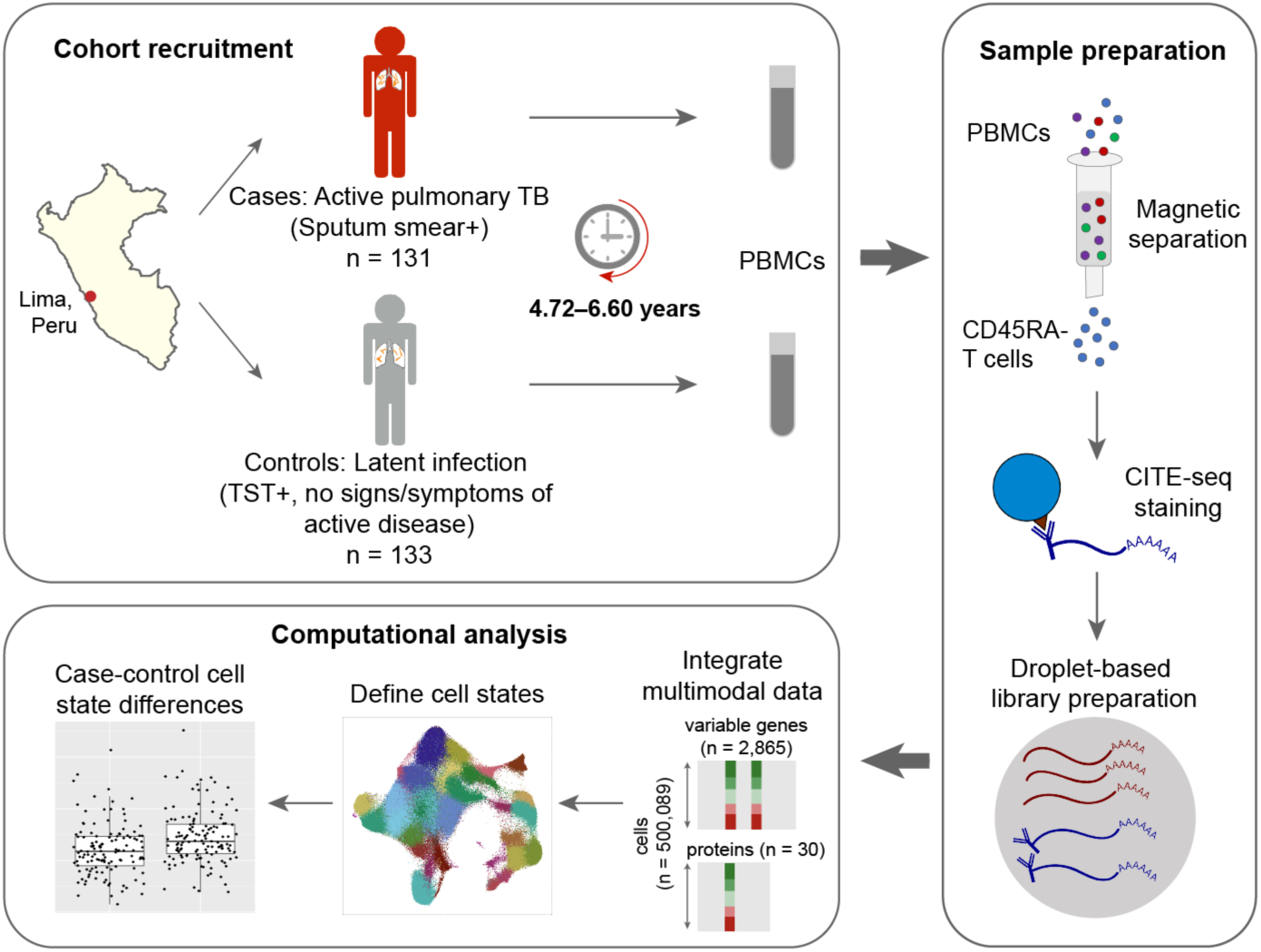
Study design. We obtained PBMCs from a Peruvian TB cohort, profiled memory T cells with CITE-seq, and integrated multimodal single-cell profiles to define cell states and case-control differences.

### Multimodal sequencing produces 500K robust T cell profiles

To profile memory T cells, we obtained peripheral blood mononuclear cells (PBMCs) from 131 cases and 133 controls, and used magnetic sorting to negatively select memory T cells at high purity (∼98.4%, **Supplementary Information, Fig. 1)**. Because T cell phenotypes can be characterized by both transcriptional and surface protein markers^27-33^, we profiled single cells with Cellular Indexing of Transcriptomes and Epitopes by Sequencing (CITE-seq), a multimodal method that combines unbiased single-cell RNA-seq with surface marker profiles obtained via oligonucleotide-tagged antibodies^34,35^. We optimized a panel of 31 surface markers (**Supplementary Table 3**), including markers of lineage (e.g., CD4, CD8), activation (e.g., CD25, HLA-DR), migration (e.g., CCR6, CXCR3), and other functions, and mouse immunoglobulin G (IgG) as a control. After cell- and sample-level quality control (**Extended Data Fig. 1a-f**), the final data set contained 500,089 memory T cells from 259 individuals (mean: 1,845 cells/sample, 95% CI: 518–3,172, **Extended Data Fig. 1g**).

As others have previously observed, mRNA-protein correlations were modest but positive, with Pearson r < 0.5 for most gene-protein pairs (**Supplementary Information**)^34,35^. To assess the accuracy of surface marker measurements with CITE-seq, we examined 8 populations gated with both CITE-seq and flow cytometry in samples from the same donor (**Supplementary Information, Supplementary Table 4**). The average frequency of each gated population was concordant between platforms (Pearson r = 0.99), and for each population, frequencies measured by the two platforms were well-correlated across individuals (Pearson r = 0.73–0.94).

### Multimodal integration defines the memory T cell landscape

To define high-resolution memory T cell states, we assume that biologically relevant states are reflected in both mRNA and surface protein signatures and can be more precisely defined by integrating both modalities. We used canonical correlation analysis (CCA) to project each cell into a low-dimensional space defined by correlated modules of transcripts and proteins (**Extended Data Fig. 2a**). This allows us to leverage signatures involving different modality-specific markers; for example, regulatory T cells have high surface expression of CD25 and absence of CD127 (IL-7R), but also express *FOXP3* transcripts^36,37^. We selected the top 20 canonical variates (CVs) with highest mRNA-protein correlations and corrected them for donor and batch effects (**Extended Data Fig. 2b, c**)^38^. The first CV is computed such that it captures the most variation shared between modalities, and accordingly, scores on this dimension correlated with a previously defined signature of cytotoxic potential (Pearson r = 0.90)^27^ (**Extended Data Fig. 2d, e**).

Applying graph-based clustering to the top 20 CVs, we defined 31 clusters representing putative cell states characterized by marker genes and surface proteins (**Fig. 2, Extended Data Fig. 3, Supplementary Table 5**). Based on protein expression, the majority of clusters (23/31) were CD4+, while 5 were CD8+. One cluster (C-24) was a mixture of CD4+ and CD8+ cells, and two (C-30 and C-31) were CD4-CD8-. Most cells in these double-negative clusters expressed *TRDC*, the constant region of the T cell receptor (TCR) delta chain, but not the alpha beta TCR surface proteins, consistent with gamma delta (γδ) T cells. Some CD4+ clusters expressed surface markers of known T cell phenotypes: CD62L marked 4 clusters as central memory^39^. Clusters C-5 and C-9 express CD25 and lack CD127 surface protein, resembling regulatory T cells^37^. Among the CD8+ clusters, we identified one central memory cluster (C-25) and distinct transitional *GZMK*+ (C-28) and cytotoxic *GZMB*+ (C-29) effector subsets^40^. Clusters C-15 and C-27 have high expression of HLA-DR and CD38 surface protein and proliferation-associated transcript *MKI67* and represent chronically activated CD4+ and CD8+ memory, respectively^41^. The observation of known cell types defined by coordinated expression of transcripts and surface markers validated our unbiased clustering approach for high-resolution dissection of the human memory T cell pool.

**Fig. 2.**
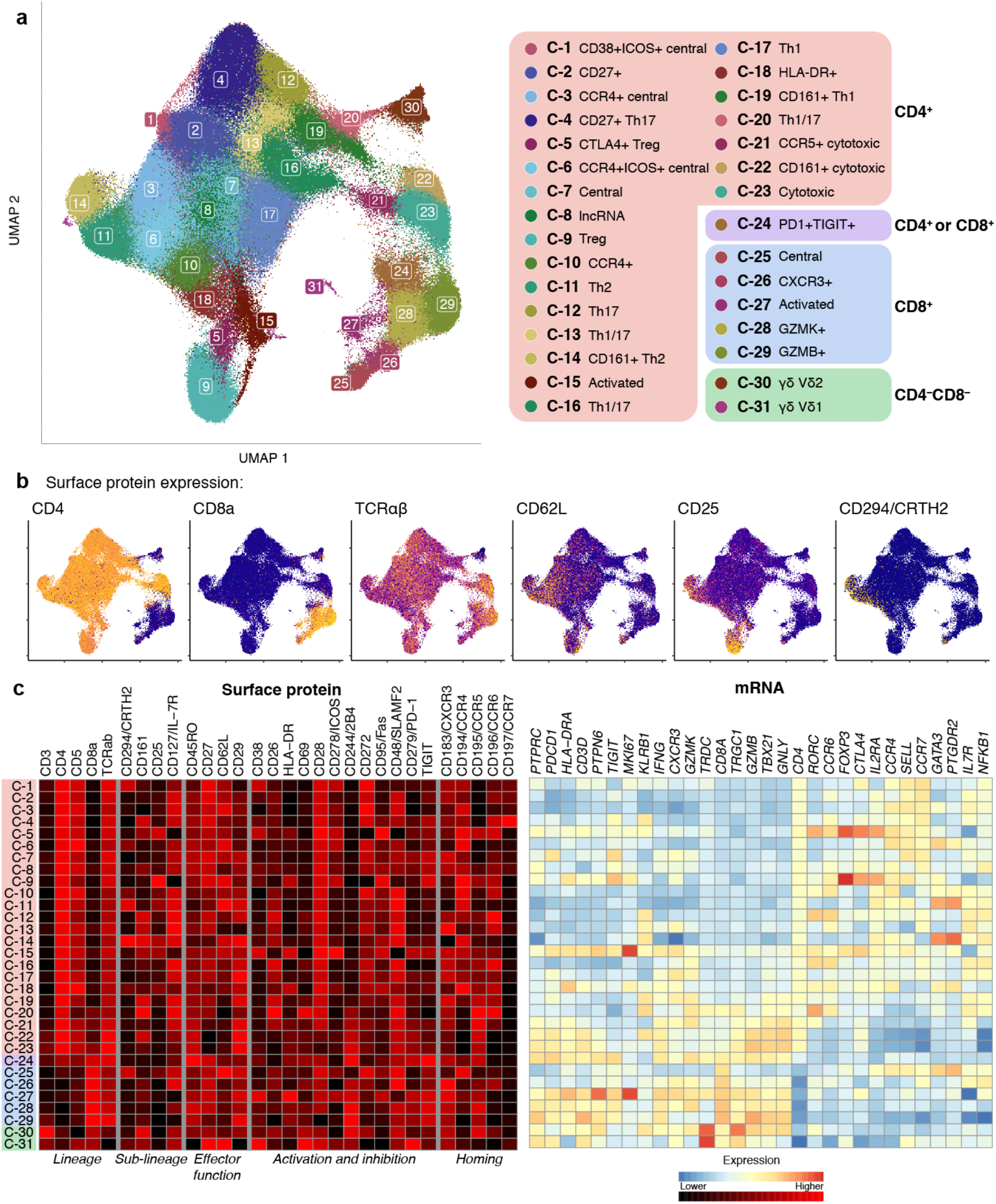
Landscape of memory T cell states. **a**, UMAP colored by 31 multimodal clusters. Cluster annotations are based on top differentially expressed genes and surface proteins. Clusters boxed in red are CD4+, purple are mixed CD4+ and CD8+, blue are CD8+, and green are CD4-CD8-. **b**, Expression of major lineage-defining surface markers measured through CITE-seq. Colors are scaled independently for each marker from minimum (blue) to maximum (yellow) expression. **c**, Heatmap of selected marker genes. Surface protein heatmap colors are uniformly scaled for each protein. mRNA heatmap colors reflect z-scores for each gene.

### Memory T cell states are associated with host factors

T cell state abundances varied widely across individuals (e.g., C-14 range: 0.07– 35% of cells per donor) but were concordant between technical replicates from the same donor (Pearson r = 0.44–1.00) (**Extended Data Fig. 4, Supplementary Information**), indicating that T cell states may be associated with other donor traits. We tested 38 demographic, socioeconomic, and genetic ancestry covariates (**Supplementary Table 6**) for association with T cell states with Mixed-effects modeling of Associations of Single Cells (MASC), a single-cell model that calculates the odds ratio of a cell being in each cluster given a covariate of interest, after correcting for other cell- and donor-level confounders^32^. We corrected for donor, batch, and total unique molecular identifiers (UMIs) and percent mitochondrial (MT) UMIs per cell, and confirmed with permutation analysis that this model obtained reliable type I error estimates (**Supplementary Information**).

Age, sex, winter blood draw, and proportion of European ancestry were significantly and independently associated with T cell state composition (**Extended Data Fig. 5**). As previous studies suggest^42^, age influenced T cell states and was the most significant covariate (**Methods**, gamma p = 2.24 × 10^−53^), with 12/31 associated states after correcting for multiple hypothesis testing (LRT p < 1.6 × 10^−3^ = 0.05/31). Similar to findings in prior reports^43,44^, cytotoxic CD4+ T cells (C-23) were expanded ∼20% per decade of age (odds ratio [OR] = 1.19 per 10 years, 95% confidence interval [CI]: 1.10–1.28, p = 1.59 × 10^−5^), while Vδ1 T cells (C-31) were reduced by more than 50% per decade (OR = 0.46 per 10 years, 95% CI: 0.39–0.54, p = 1.83 × 10^−22^, **Extended Data Fig. 6a, Supplementary Table 6**). Sex was also strongly associated with T cell states (gamma p-value = 8.40 × 10^−28^). For example, *GZMB*+ CD8+ T cells were expanded in males (C-29: OR = 1.88 M vs. F, 95% CI: 1.46–2.42, p = 5.51 × 10^−5^), and Th1s were expanded in females (e.g., C-17: OR = 0.77 M vs. F, 95% CI: 0.71–0.83, p = 1.20 × 10^−11^) (**Extended Data Fig. 6a, Supplementary Table 6**), recapitulating previously published observations^45,46^.

We also observed surprising associations of T cell states to season of blood draw (winter gamma p = 4.40 × 10^−24^). Th2 states were expanded in samples drawn in the winter (e.g., C-11: OR = 1.24, 95% CI: 1.10–1.39, p = 5.13 × 10^−4^) (**Extended Data Fig. 6a, Supplementary Table 6**). To our knowledge, this has not been reported previously, although studies have noted the seasonality of cytokine responses^47^. Immune function has also been shown to vary with ancestry^48^, and in our cohort, individuals with higher European ancestry showed nominal depletion of all three clusters of cytotoxic CD4+ T cells (European gamma p = 2.21 × 10^−5^, e.g., C-23: OR = 0.14 per percent European ancestry, 95% CI: 0.05–0.48, p = 0.03) (**Extended Data Fig. 6a, Supplementary Table 6**). Notably, age and sex were also associated with TB disease progression (**Supplementary Table 1**), but along with season of blood draw and ancestry, maintained associations with T cell state abundances even after adjusting for TB progression status (**Extended Data Fig. 6b, Supplementary Table 7**) and in a joint model (**Extended Data Fig. 6c, Supplementary Table 8)**.

### An *RORC*-expressing effector state is reduced in TB progressors

We used a MASC model to identify T cell states associated with TB disease progression, after adjusting for potentially confounding covariates (age, sex, winter blood draw, and proportion of European ancestry) and batch and single-cell technical factors (**Methods**). We observed a significant reduction in cluster C-12 in individuals with a history of TB disease after correction for multiple hypothesis testing (OR = 0.80, 95% CI: 0.73–0.87, p = 1.21 × 10^−6^) (**Fig. 3a, Supplementary Table 8**). Notably, C-12 was also independently associated with other covariates in the full model, demonstrating depletion with increased age (OR = 0.82, p = 2.69 × 10^−3^) and in males (OR = 0.85, p = 4.30 × 10^−4^), and expansion in winter blood draws (OR = 1.16, p = 1.30 × 10^−3^, **Extended Data Fig. 6d**). This covariate-aware multimodal approach is well-powered to detect even a modest case-control difference in C-12 frequency (mean: 3.0% in cases, and 3.6% in controls, **Fig. 3b**), which may have been obscured with unimodal clustering failing to precisely capture this cell state (**Extended Data Fig. 7, Supplementary Table 9** and **10**).

**Fig. 3.**
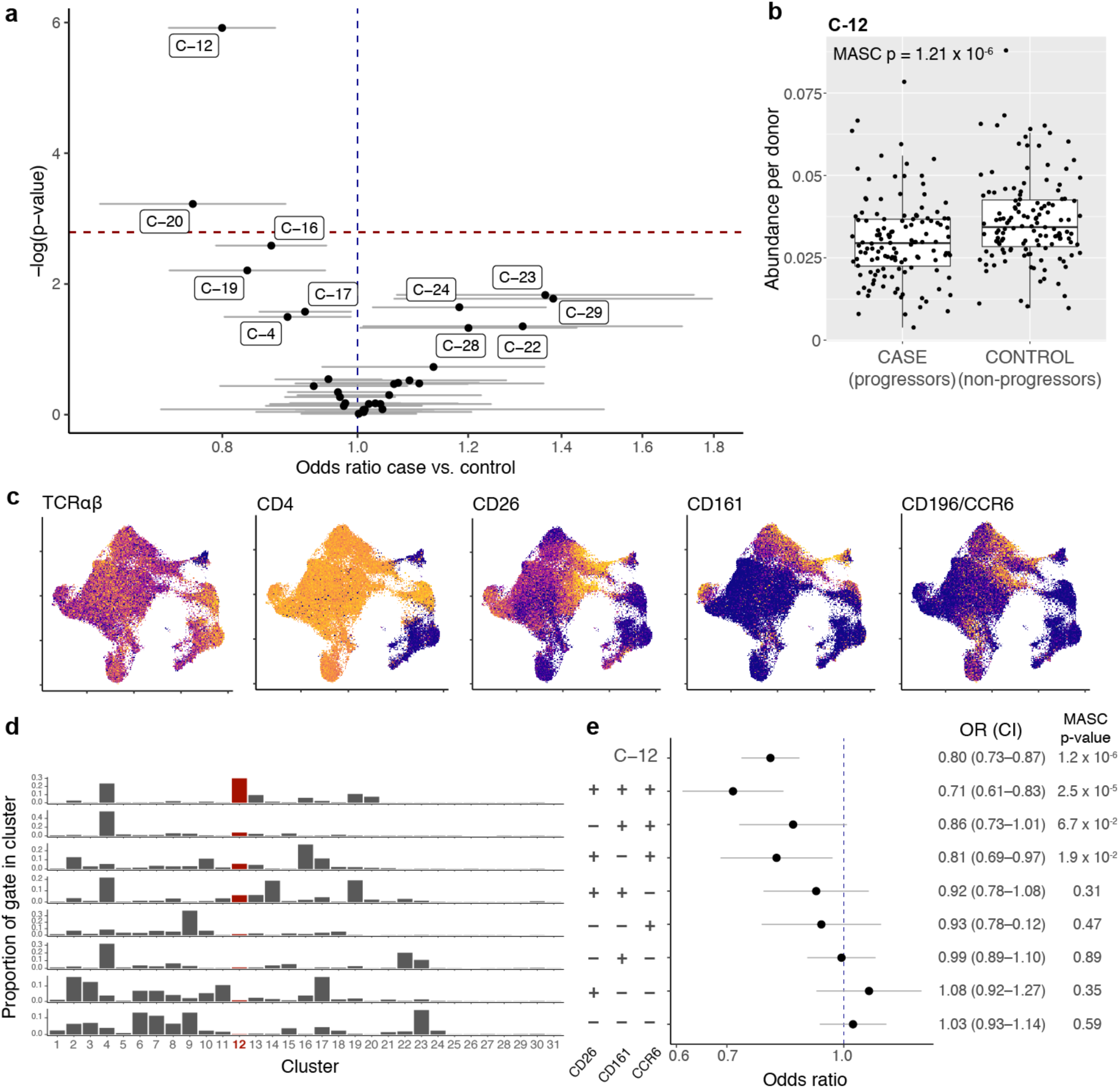
Identification and isolation of a depleted memory T cell state in TB cases. **a**, Associations between TB disease status and memory T cell states. Each multimodal cluster is plotted based on the MASC odds ratio of a cell being in that cluster for cases vs. controls, and the -log(LRT p-value) of the association. Error bars show the 95% confidence interval. The red dashed horizontal line corresponds to a Bonferroni-adjusted p-value threshold of 0.05/31 = 1.6 × 10^−3^. Labeled clusters are significant at a nominal p-value threshold of p < 0.05. **b**, Abundance of C-12 in 128 cases and 131 controls. **c**, Surface protein markers of C-12. Colors are scaled independently for each marker from minimum (blue) to maximum (yellow) expression. Boxplot center line = median, box limits = 25^th^ and 75^th^ percentile, whiskers = 1.5x interquartile range. **d**, Distribution of each gate across clusters. **e**, Association of gated populations with TB progression status. For each gated population, we plotted the MASC odds ratio of a cell having that phenotype for cases vs. controls. P-values are from an LRT with 1 degree of freedom (d.f.). Error bars show the 95% confidence interval.

Cells in this cluster have a CD4+ effector surface phenotype (CD62L: Expression fold change in vs. out of C-12 [FC] = 0.66, p = 2.99 × 10^−7^, CCR7: FC = 0.85, p = 9.72 × 10^−3^) and lack surface markers for activation (CD38: FC = 0.56, p = 2.30 × 10^−10^, HLA-DR: FC = 0.39, p = 1.86 × 10^−18^) or exhaustion (PD-1: FC = 0.76, p = 7.23 × 10^−6^, TIGIT: FC = 0.26, p = 1.10 × 10^−33^). The top surface protein markers were CD26, CCR6, and CD161 (**Fig 3c, Supplementary Table 5**) and the top mRNA markers were *CCR6, CTSH*, and *KLRB1*, with elevated expression of transcripts for Th17 lineage-defining transcription factor *RORC* (FC = 5.70, p = 2.37 × 10^−187^) and reduced expression of transcripts for Th1 lineage-defining *TBX21* (FC = 0.52, p = 1.01 × 10^−10^) and *IFNG* (FC = 0.30, p = 7.23 × 10^−34^). This combination of surface protein and mRNA markers suggests that C-12 represents a subset of Th17 cells^49,50^. The C-12 state was not changed by TB disease status; there were no differentially expressed genes between the cases and controls in this cluster.

We also observed less significant reduction of cluster C-20 in cases (OR = 0.76, 95% CI: 0.65–0.89, p = 5.95 × 10^−4^, **Extended Data Fig. 8)**. C-20 contained an average of 1.1% and 1.2% of cells in cases and controls, respectively, and expressed CD26, CD161, CCR6, and CCR5 surface markers and *DPP4, KLRB1*, and *BHLHE40* mRNA markers. Transcripts for Th1 lineage-defining transcription factor *TBX21* were also highly expressed (FC = 1.70, p = 1.65 × 10^−7^), as well as Th17 lineage-defining transcription factor *RORC* (FC = 5.11, p = 2.98 × 10^−160^). These findings suggest that C-20 is distinct from C-12 and has a mixed Th1/17 phenotype^51^.

Because our cohort was profiled several years after TB disease was diagnosed and treated, we did not expect differences in activated cell states. Using MASC, we verified that there were no differences in CD4+ HLA-DR memory T cells (OR = 1.02, CI = 0.86–1.22, p = 0.79, **Supplementary Table 11**). Additionally, because we profiled all memory T cells, it is unsurprising that other differences implicated in previous TB progression studies of *M*.*tb* antigen-specific cells were only weakly or marginally significant (**Supplementary Table 11**).

### C-12 matches a CD26+CD161+CCR6+ phenotype

One advantage of multimodal profiling is that it enables targeted validation. To functionally characterize C-12 (the most significant association with TB disease progression), we used surface protein measurements from CITE-seq to identify sortable markers. We built a classification tree to identify a minimal set of candidate markers for C-12— CD26+, CD161+, and CCR6+—and confirmed with stepwise backward selection that removing any one of these markers reduced our sensitivity for C-12 (sensitivity = 54.8%, specificity = 95.5%, **Fig. 3d, Methods, Supplementary Information**). If this population corresponds to the C-12 state, it should be associated with TB progression when gated in CITE-seq data. Indeed, MASC modeling demonstrated reduction in TB cases (OR = 0.71, CI = 0.61–0.83, p-value = 2.47 × 10^−6^, **Fig. 3e**). Removing any individual gate weakens the association.

### C-12 produces IL-17 and IL-22

To define the cytokine profile of C-12, we isolated CD4+ T cells from three Boston-based donors who had not been assessed for TB infection and sorted CD45RO+CD26+CD161+CCR6+ cells (**Supplementary Information, Supplementary Table 12**). For comparison, we sorted naive CD4+, other memory CD4+, and regulatory T cells (**Methods**). We stimulated each population with CD3/CD28 beads for pan-TCR activation and measured a broad range of T helper cytokines in the supernatant. Compared to all other memory CD4+ T cells, our target population had higher expression of IL-17A, IL-17F, and IL-22 and lower expression of IL-4 and IL-13 (two-sided t test, IL-17A: t = 5.07, p = 0.04; IL-17F: t = 6.34, p = 0.02; IL-22: t = 8.00, p = 0.012; IL-4: t = -6.96 p = 0.02; IL-13: t = -4.44, p = 0.02, **Fig. 4a**). To determine if this observation was robust to stimulation condition and assay, we stimulated cells from five Boston donors with phorbol 12-myristate 13-acetate (PMA) and ionomycin, and with intracellular staining, again found that our target population was more likely to produce IL-17A, IL-17F, and IL-22 than other CD4+ memory T cells (Cochran-Mantel-Haenszel [CMH] OR, IL-17A: 12.6, IL-17F: 18.1, IL-22: 6.0, all p < 0.001, **Fig. 4b-d, Extended Data Fig. 9**). Inverting any one of the three surface markers (CD26, CD161, and CCR6) reduced the proportions of IL-17A, IL-17F, and IL-22 expression. This population was also polyfunctional and produced IFNγ and TNF at similar rates to CD4+ memory T cells overall (CMH OR, IFNγ: 0.72, TNF: 1.59, all p < 0.001). Notably, the target Th17 subset (CD26+CD161+CCR6+) had more than three times as many IFNγ-producing cells as the rest of the Th17 compartment (two-sided t test vs. CD26-CD161+CCR6+, p = 2.23 × 10^−5^, **Extended Data Fig. 9a**).

**Fig. 4.**
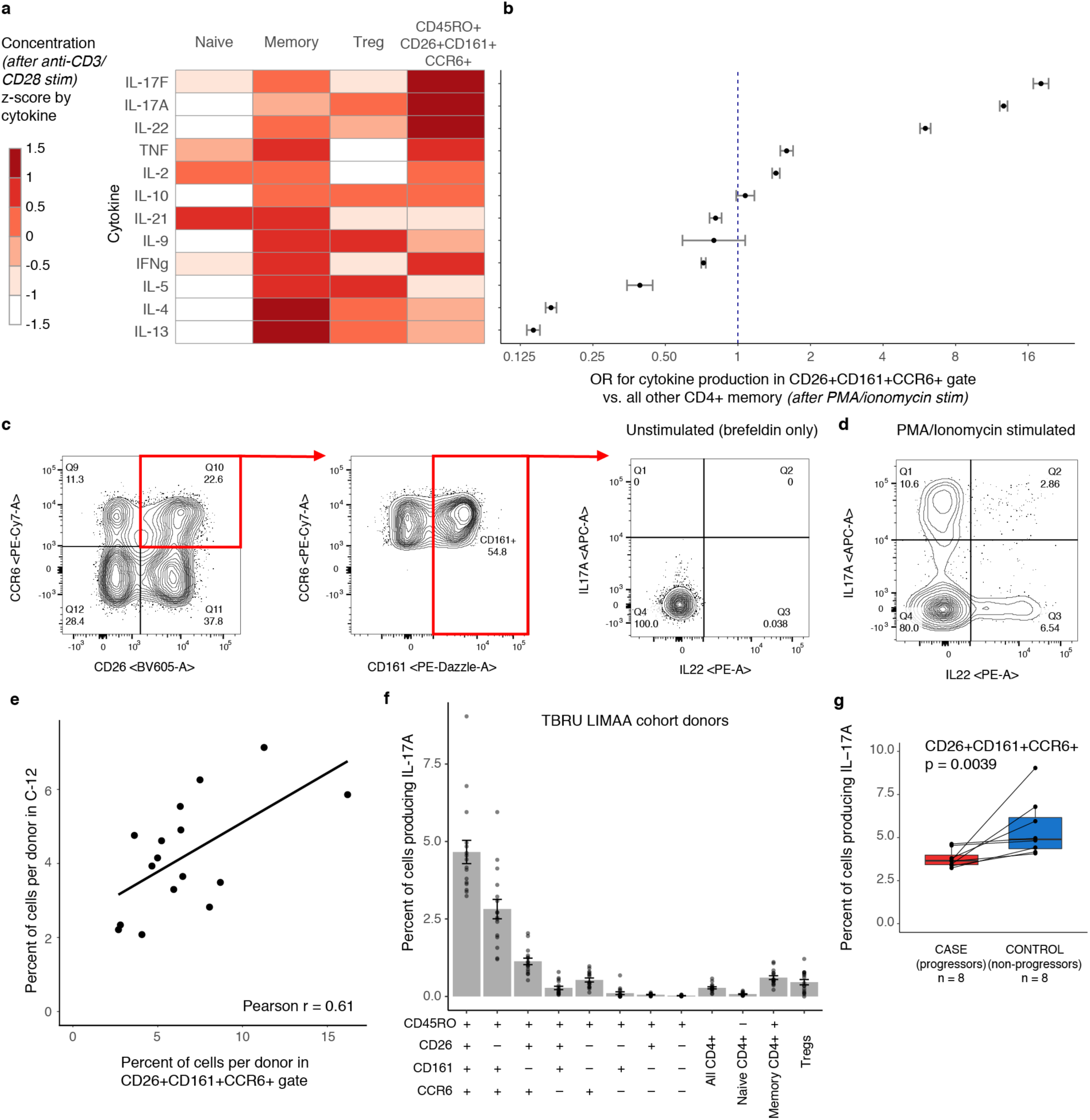
Characterizing C-12 as an IL-17+ state with reduced function in Peruvian TB cases. **a**, Cytokine expression in supernatant for four CD4+ T cell subsets after CD3/CD28 bead stimulation. We estimated cytokine concentrations and averaged across donors (n = 3), scaled by cytokine, and binned into sextiles. **b**, Odds ratio of cytokine expression in gated population based on ICS (n = 5). We used the Cochran-Mantel-Haenszel method to calculate the odds ratio of cytokine production inside vs. outside the gate. Bars show the 95% confidence interval. **c**, Gating strategy to isolate CD26+CD161+CCR6+ memory CD4+ T cells. Intracellular staining for IL-17A and IL-22 is shown without stimulation. **d**, Intracellular staining for IL-17A and IL-22 after stimulation with PMA/ionomycin. **e**, Correlation between abundance of flow-gated population and C-12, per donor. We calculated a linear best fit and the Pearson correlation coefficient. **f**, Per-donor percent of cells producing IL-17A in populations gated in Peruvian TB cohort donors. Bars represent the mean and error bars show standard error of the mean across 16 donors. **g**, Case-control comparison of per-donor percent of cells in gated population producing IL-17A. Paired samples were matched for age, sex, season of blood draw, and proportion of European ancestry. We calculated p-values from a one-sided Wilcoxon signed-rank test (test statistic V for IL-17A = 0). Boxplot center line = median, box limits = 25^th^ and 75^th^ percentile, whiskers = 1.5x interquartile range.

Because IL-17 and IL-22 best characterized this population’s functional phenotype in disease-unascertained non-Peruvian donors, we next measured these cytokines in Peruvian donors with a history of *M*.*tb* infection. We selected eight pairs of cases and controls from our original CITE-seq cohort, matched for age (+/-5 years), sex, season of blood draw, and proportion of European ancestry (+/-0.05); isolated CD4+ T cells and stimulated with PMA and ionomycin; and measured IL-17A and IL-22 production in target (CD45RO+CD26+CD161+CCR6+) and control populations. The frequency of the target population correlated well with the per-donor abundance of cluster C-12 (r = 0.61, **Fig. 4e**) and with *in silico*-gated proportions (r = 0.62, **Extended Data Fig. 10a**) in our CITE-seq data.

Similar to Boston samples, in Peruvians this population contained more cells that produce IL-17A (mean = 4.7 ± 1.5% of cells in gate) or IL-22 (mean = 2.9 ± 2.0%) in response to stimulation than any other Boolean combination of the three gates (**Fig. 4f, Extended Data Fig. 10b**). In fact, despite making up on average only 6.6% of CD4+ memory T cells, the target population comprised over one-third of total IL-17A or IL-22-producing CD4+ memory cells (IL-17: 49.2%, IL-22: 33.2%) (**Extended Data Fig. 10c**), and were 14.3 times as likely to produce IL-17A and 6.7 times as likely to produce IL-22 (CMH OR, p < 0.001, **Extended Data Fig. 10d**).

### C-12 produces less IL-17 in people with a history of TB disease

We hypothesized that in individuals who progress to TB disease, C-12 may not only be reduced in frequency, but in function as well. To test this, we compared cytokine production in our target population between matched cases and controls. We observed that, in response to PMA and ionomycin stimulation, only 3.8% of CD4+CD26+CD161+CCR6+ memory T cells in cases produced IL-17A, compared to 5.5% in controls (one-sided paired Wilcoxon signed rank test p = 0.0039, **Fig. 4g**). This functional deficiency was specific to our target population, compared to broader populations, including CD4+ memory T cells overall (p = 0.097, **Extended Data Fig. 10e**). IL-22 production was also lower in cases than controls in CD4+CD26+CD161+CCR6+ memory cells, but with weaker difference (p=0.025, **Extended Data Fig. 10e**).

## Discussion

In this study, we survey the full spectrum of memory T cells in TB cases and infected controls outside the context of active disease, and show that the most significant steady-state differences reside in a relatively rare (∼3% of memory T cells) Th17 subset (C-12) marked by a CD4+CD45RO+CD26+CD161+CCR6+ surface phenotype, production of IL-17 and IL-22 upon stimulation, and higher IFNγ production than other Th17s. This cell state is not only less abundant in progressors but also produces significantly less IL-17 and IL-22.

We observe both of these differences in patients who recovered from TB and without restricting analysis to antigen-specific cells, which suggests that observed differences in C-12’s abundance and functionality may (1) be a long-term consequence of prior TB disease, or (2) predispose individuals to TB disease progression. This latter hypothesis is plausible, first, because T cell phenotypes previously identified in antigen-specific cells during TB disease share aspects of the C-12 phenotype. Human immunoprofiling studies have shown that *M*.*tb* antigen-responsive cells produce less IL-17 and IL-22 in TB cases compared to healthy controls^19^, and are reduced in progressors during disease compared to latency^21^. The bulk of *M*.*tb* antigen-specific cells have been mapped to a Th1/17 polyfunctional CCR6+CXCR3+ state that is expanded in latent individuals compared to progressors during active disease^22,49,52^. Although studies have emphasized that CCR6+CXCR3+ cells produce IFNγ upon *M*.*tb* antigen stimulation, in non-human primates this cell state also produces IL-17 in bronchoalveolar lavage and is expanded in the lungs during latent infection compared to TB disease^53^. Second, mutations in *RORC*—the Th17 lineage-defining transcription factor that is highly expressed in C-12—increase susceptibility to mycobacterial diseases^5^. Third, C-12 is also depleted with increased age, in males, and outside of winter (**Extended Data Fig. 6d**), all of which have been associated with elevated TB risk^2,54^. These host variables may in part increase TB risk by reducing C-12 frequencies. Finally, C-12 also does not have an activated or exhausted phenotype, arguing that they are not the consequence of chronic stimulation.

Prospective studies profiling the immune system prior to infection are required in order to conclusively establish a causal link. Our results demonstrate the power of high-dimensional T cell profiling in human cohorts to multimodally define novel cell states and identify steady-state differences in the immune system that underlie divergent disease outcomes.

## Methods

### Clinical cohort

Our cohort is a selected subset of 14,044 individuals from a large epidemiological parent study of risk factors for TB infection and disease conducted between 2008 and 2012 (protocol approved by the Harvard University Institutional Review Board [#19332] and by the Research Ethics Committee of the National Institute of Health of Peru)^25^. All study participants were recruited from 106 district health centers in Lima, Peru and provided written informed consent. We enrolled index patients aged 16 or older with microbiologically confirmed pulmonary TB. Within two weeks of enrolling an index patient, we enrolled their household contacts who were assessed for co-prevalent TB disease by clinical evaluation and for TB infection by a tuberculin skin test (TST). Household contacts were reassessed at 2, 6, and 12 months for evidence of new TB infection or TB disease.

At the time of enrollment, we collected demographic, health, and socioeconomic data from both index patients and household contacts, including age, sex, height, weight, alcohol use, smoking, prior incarceration, Bacillus Calmette-Guérin (BCG) vaccination scars, isoniazid preventative therapy, and previous TB diagnosis. Nutritional status was determined for children based on the World Health Organization BMI z-score tables, and for adults based on BMI thresholds. Individuals were categorized as underweight (children, age ≤ 19: z-score ≤ -2; adults: BMI < 18.5), normal weight (children: -2 < z-score ≤ 2; adults: 18.5 ≤ BMI < 25), or overweight (children: z-score > 2; adults: BMI ≥ 25). Alcohol use was categorized as non-drinker (0 alcoholic drinks per day), light (< 40 grams or < 3 alcoholic drinks per day), or heavy (≥ 40 grams or ≥ 3 alcoholic drinks per day). Smoking was categorized as non-smoker (0 cigarettes per day), light (1 cigarette per day), or heavy (> 1 cigarette per day). BCG vaccination status was self-reported. Number of BCG scars was based on physician’s observation. Socioeconomic status (SES) was categorized into tertiles based on a principal component analysis (PCA) that included type of housing, access to a water supply, and sanitation. Season of blood draw was classified based on local temperatures: winter (June-September), spring (October-December), and summer (January-May)

For this study, we re-consented and enrolled a subset of 264 participants from the parent study whom we visited to obtain information on their TB history subsequent to the completion of the parent study and to obtain PBMCs. We considered index patients and household contacts who develop TB disease during follow up as cases, and TST-positive household contacts who did not develop TB disease as controls. We excluded participants if they did not consent to re-enrollment or were HIV-positive. We collected blood a median of 5.7 years after enrollment in the parent study (range 4.72– 6.60). All cases had been infected with drug-sensitive strains, and received treatment before re-recruitment. Controls were excluded if they were first-degree relatives of their index patient.

We calculated associations between each covariate and TB disease status with either a two-sided t test (continuous covariates) or a Chi-squared test (categorical covariates). For significantly associated covariates, we re-estimated associations in a multivariate logistic regression model, and determined significance based on coefficient p-values.

### Sample Processing

#### PBMC Sample Preparation

Within 6–8 hours of obtaining blood samples, we purified PBMCs using Ficoll-Hypaque as described^55^, followed by cryopreservation at a concentration of 5 million cells/mL for shipping to Boston.

We quickly thawed cryopreserved PBMCs (10 million cells) and added each sample dropwise to pre-warmed complete RPMI (cRPMI) (RPMI 1640 supplemented with 10% heat inactivated fetal bovine serum (Gemini), nonessential amino acids (Gibco), 2-mercaptoethanol (Gibco), penicillin/streptomycin (Gibco), L-glutamine (Gibco)). We washed and resuspended cells in 1 mL of cRPMI, and saved an aliquot of each sample (5% of total cells) at 4° C for flow cytometry staining.

#### Flow Cytometry

We processed 12 samples per day on 23 days over a 15-week period. In total, we collected flow cytometric data on 276 samples (264 unique donors, with 12 technical replicates run in separate batches on consecutive weeks). We washed PBMCs in PBS and stained with blue fluorescent Live/Dead fixable dead cell stain (1:1000) (Invitrogen). We covered each sample in foil, and incubated for 20 minutes at room temperature. After centrifugation, we stained samples with an antibody master mix (**Supplementary Table 4)** in Brilliant Stain buffer (BD Bioscience, Cat #566349). We covered in foil and incubated for 25 min at 4° C. We washed samples with MACS buffer (pH 7.4 PBS, 2mM EDTA, 2% FBS) and filtered through 40um mesh prior to flow cytometry using a BD LSRFortessa™ and analysis with FlowJo Version 10.6.2. We used the gating strategy shown in **Supplementary Information** to identify lymphocytes and memory subpopulations.

#### CITE-seq

We applied an optimized version of CITE-seq to memory T cells from 276 samples (264 unique donors, with 12 technical replicates)^35^. We processed 12 samples per day on 23 days over a 15-week period. We ran technical replicates in separate batches on consecutive weeks.

To isolate memory T cells, we modified the Pan T cell negative Isolation magnetic-activated cell sorting (MACS^R^) kit (Miltenyi Biotec, Cat #130-096-535) by adding anti-CD45RA biotin (Miltenyi Biotec, Clone REA1047, 2 uL per stain) to the antibody cocktail. For 10 million cells (our expected input), we used 2x reagents to achieve memory T cell purity of 98.4% (**Supplementary Information**). After isolating memory T cells, we stained up to 300,000 memory T cells per donor. Then we centrifuged and stained each sample with FcX True Stain (BioLegend) with 0.2 ug/uL dextran sulfate sodium (Sigma-Aldrich, Cat #RES2029D-A707X) in labeling buffer (PBS 7.4 with 1% UltraPure Bovine Serum Albumin (BSA)) for 10 min at 4° C.

We then added a TotalSeq™-A (BioLegend) oligonucleotide-labeled antibody mix (anti-CCR6 suspended in 10 uL of labeling buffer) and incubated all samples at room temperature for 25 min. Next, we stained with the remaining 30 TotalSeq™-A antibodies (**Supplementary Table 3**) for 25 min at 4° C and washed cells three times with 2 mL, 1 mL, and 1mL of labeling buffer sequentially. Each sample was passed through a 40um filter and kept on ice prior to sorting on a BD FACSAria™ Fusion cell sorter.

We sorted up to 10,000 live cells from each sample based on forward and side scatter gating to remove non-lymphocytes, dead cells, and other impurities. We then pooled cells into batches of six donors. Batch assignments were randomized while requiring that no two donors in the same batch had a relatedness estimation in admixed populations (REAP) kinship score greater than 0.125 (at most, second cousins) based on genotype, to facilitate *post hoc* demultiplexing^56^. Pools of 6 samples were sorted into one Eppendorf tube prepared with 200 uL of 0.4% BSA in PBS, and each pool was processed as one scRNA-seq sample.

We prepared mRNA and surface marker libraries for each batch at the Brigham and Women’s Hospital Single Cell Genomics Core using the Chromium Single Cell 3’ v3 kit (10x Genomics). Pairs of libraries prepared on the same day were pooled and sequenced to a depth of 400 million reads per lane on an Illumina HiSeq X with paired-end 150 base-pair reads. In total, we sequenced 276 samples across 46 pooled libraries in 6 sequencing runs.

#### Genotyping and genetic data processing

We genotyped all individuals on the LIMAArray, a previously described custom Affymetrix array designed based on whole-exome sequencing from 116 Peruvian individuals with active TB^8^. Genotypes were called for all 4002 individuals in the original genetic study with the apt-genotype-axiom program. We excluded individuals with high genotype missingness (≥ 5% of loci) or high heterozygosity rate (±3 standard deviations). We excluded loci with significant association with batch (p < 1 × 10^−5^), low call rate (< 95%), large difference in per-single nucleotide polymorphism (SNP) missingness rate between cases and controls (> 10^−5^), Hardy-Weinberg (HWE) p-value below 10^−5^ in controls, and duplicated position markers. After individual and SNP-level quality control, there were 263 donors and 677,385 SNPs remaining.

To measure global ancestry proportions for each donor, we joined our cohort with previously published genotypes from the 1000 Genomes Project phase 3 (2,054 individuals from 26 populations)^57^, Siberians (245 individuals from 17 populations, and Native Americans (493 individuals from 57 populations) based on variant-level matching^58^. After removing variants with minor allele frequency (MAF) < 1%, 34,936 variants remained. We performed PCA and pruned for linkage disequilibrium (LD, r^2^ > 0.1 between any pair of markers within a sliding window of 50 markers with 10-marker offset) with PLINK (version 1.90b3w). We used the 22,266 remaining variants to measure global ancestry with ADMIXTURE (version 1.3) at K = 4^59,60^. Because of the admixed nature of the cohort, we calculated an admixture-aware genetic relatedness matrix with the REAP kinship score to account for linkage disequilibrium differences^56^.

We pre-phased genotypes with SHAPEIT2 and imputed genotypes at untyped autosomal loci with IMPUTE2, using the 1000 Genomes Project Phase 3 dataset as a reference panel^61,62^. After removing SNPs with INFO scores less than 1, 738,194 SNPs remained.

### Statistical Analysis of Genomic Data

#### Quantifying surface markers and genes

We used Cell Ranger (version 3.1.0) to conduct all alignment and feature quantification of multimodal single-cell sequencing data. For mRNA, we aligned reads to the human genome (GRCh38 for transcriptomic analysis and hg19 for genotype-based demultiplexing). We aligned surface protein reads to a dictionary of feature tags. We collapsed reads mapping to the same gene or surface marker in the same cell to a single unique molecular identifier (UMI).

#### Single-cell sample demultiplexing

We demultiplexed the six samples within each pooled batch based on genotypes at 738,194 SNPs. We used Demuxlet with default parameters and removed cells with ambiguous or doublet assignments^63^, and verified the accuracy by correlating the number of cells demultiplexed per sample with the number of live cells sorted after memory T cell isolation.

#### Single-cell sequencing data quality control

We removed cells that expressed fewer than 500 genes or had more than 20% of their UMIs mapping to MT genes (**Extended Data Fig. 1a, b**). Gene expression UMI counts were normalized per cell for library size and log-transformed: ln 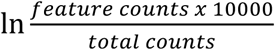

For samples with high live cell counts but low numbers of demultiplexed cells, we merged single-cell sequencing reads assigned to each donor, and called variants from merged data using bcftools (v1.9)^64^. We quantified the concordance between sequencing-based genotypes and array-based genotypes and corrected the donor labels for 4 samples. We identified and removed an additional 4 mislabeled samples. One more sample was removed for high genotype missingness and high heterozygosity rate.

We normalized the surface marker UMI counts using a centered log ratio transformation for each cell. We used an *in silico* gating strategy to identify and remove contaminating non-memory T cells with low CD3 and/or low CD45RO surface marker expression. We biaxially plotted cells based on their normalized expression of CD3 and CD45RO, manually determined thresholds of each marker’s expression to separate discrete subpopulations, and removed cells with expression of either marker below those thresholds (**Extended Data Fig. 1c-e**). We additionally removed cells that were in clusters dominated by non-memory T cells.

#### Unimodal pipeline for dimensionality reduction

For each modality, we selected the union of the top 1000 features with highest variance in each library preparation pool, and scaled the expression of each feature across all cells to have mean = 0 and variance = 1. For the mRNA expression, we also cosine normalized the scaled expression values. We used PCA to reduce the data into 20 dimensions, and then corrected these PCs for donor and library preparation batch effects using Harmony^38^. With uniform manifold approximation and projection (UMAP), we reduced the batch-corrected embeddings into two dimensions for improved visualization^65^.

#### Multimodal pipeline for dimensionality reduction

For each modality, we selected the union of the top 1000 features with highest variance in each library preparation pool, and scaled the normalized expression of each feature across all cells to have mean = 0 and variance = 1. We excluded T cell receptor genes because of potential mapping errors due to recombination and sequence similarity. Then, we used CCA as implemented in the cc function from the *CCA* R package to calculate canonical dimensions defined by correlated gene and protein expression^66^ (**Extended Data Fig. 2a**). This method finds maximally correlated linear combinations of features from each modality, i.e., calculates vectors *a* and *b* for mRNA matrix *X* and surface protein matrix *Y* to maximize *cor*(*Xa, Yb*) subject to the constraint that *var* (*Xa*) = *var*(*Yb*) = 1. We defined canonical variates by projecting cells onto each canonical dimension in the mRNA space (*CV*1 = *Xa*_1_) and selecting the top 20 dimensions defined by highest canonical correlations *cor*(*Xa*_*i*_, *Yb*_*j*_) between mRNA and protein. We corrected donor and batch effects with Harmony and reduced the batch-corrected embeddings into two dimensions with UMAP.

#### Clustering and annotating cell states

Cells were clustered based on their low dimensional embeddings (either PCs or CVs). We constructed a shared nearest neighbor graph and conducted Louvain modularity clustering at a range of resolutions. Results are shown at a resolution of 2.00, which yielded 31 CCA-based clusters with at least 10 cells from more than five donors.

To annotate clusters as cell states, we identified differentially expressed mRNA and surface protein features between cells inside and outside of each cluster. Because single-cell mRNA features are sparse, we collapsed single-cell expression profiles for each modality into pseudo-bulk profiles by summing the raw UMI counts for each gene or surface protein across all cells from the same donor, batch, and cluster. For mRNA, we limited differential expression analysis to genes that had at least 30 UMIs detected in at least 120 pseudo-bulk samples (n = 4,540), and for both modalities, we normalized counts for each feature in each pseudo-bulk sample into counts per million (CPM). We used linear models to model the effect of each cluster individually on pseudo-bulk expression of each gene, accounting for donor, batch, and the number of UMIs assigned to each pseudo-bulk sample. P-values were obtained through a likelihood ratio test (LRT) between the models with and without the cluster term. We considered a gene or surface protein to be a marker of a cluster if it had a p-value < 0.05/(4,540 genes x 31 clusters) = 3.6 × 10^−7^ and a fold-change > 2. We manually annotated each cell state based on literature about its markers.

### Testing cell populations for association with TB disease progression

We tested the association of each cell state with TB disease status with MASC^32^. We specified the number of UMIs and percent MT UMIs as cell-level covariates, and donor and library preparation batch as random effects, and fit the following logistic model for each cluster *j*:

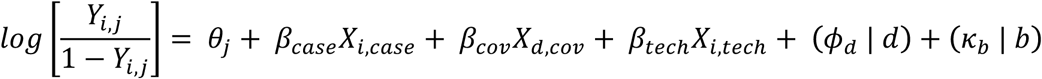

where *Y*_*i,j*_ is the odds of cell *i* being in cluster *j, θ*_*j*_ is the intercept for cluster *j, β*_case_ is the effect estimate (log(OR)) for case-control status, *β*_cov_ is a vector of effect estimates for each donor-level covariate, *β*_tech_ is a vector of effect estimates for each technical cell-level covariate, and *X*s are the corresponding values for either cell *i* or donor *d*, as appropriate. *ϕ*_d_ | *d* is a random effect for cell *i* from donor *d*, and *κ*_b_ | *b* is a random effect for cell *i* from batch *b*.

With stepwise forward selection, we identified donor-level covariates that significantly influence cell state abundance. We used MASC to test for differentially abundant clusters associated with each covariate individually, after correcting for the cell-level and batch covariates. For each covariate, this test yielded 31 cluster-specific LRT p-values. Under a null hypothesis, where the covariate is not significantly associated with cell state abundance, these cluster p-values should follow a gamma distribution parametrized by rate = 31 (the number of clusters) and scale = 1. We calculated a gamma test statistic and p-value quantifying how much the distribution of p-values for each of the 38 donor-level covariates deviates from this null, and selected the most significant covariate to add to the model. Then, we repeated this process for each remaining covariate with the expanded model, for a total of 5 iterations.

Based on this model selection process, we specified age, sex, winter blood draw, and percentage of European ancestry as donor-level fixed effects, number of UMIs and percent MT UMIs as cell-level fixed effects, and donor and library preparation batch as random effects in our full MASC model:

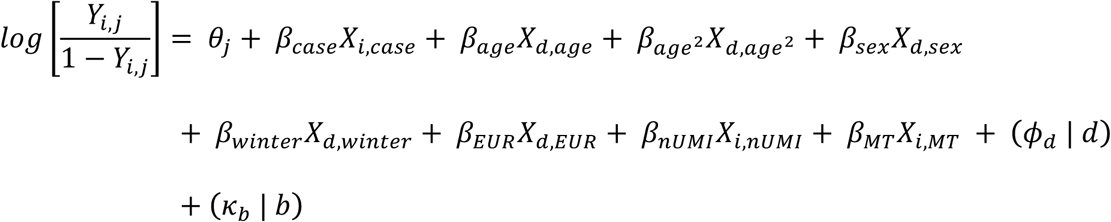

Age was included as both a linear and a quadratic effect to capture non-linear effects of age. We used Bonferroni correction to account for multiple hypothesis testing. To verify that MASC has a well-controlled type 1 error, we ran 1,000 trials of a MASC model testing the association of cluster abundances with permuted TB disease status, adjusting for age, sex, donor, batch, percent MT UMIs per cell and number of UMIs per cell. We binned percent MT UMIs and number of UMIs into quintiles. To measure case-control associations with populations from previous TB disease progression studies (**Supplementary Table 11**), we gated these populations based on normalized surface protein expression measured in CITE-seq, and used the same MASC model specified above to estimate the OR.

### Defining surface marker gates *in silico*

We used the single-cell surface marker expression to define flow-cytometry gates to isolate the CCA-defined cluster shown to be expanded in latent controls. We started with TCRαβ+, CD4+, and CD8-gates based on observed protein markers in the CITE-seq data (**Fig. 2b, Supplementary Table 5**). Then, using a classification and regression tree (CART) model implemented in R with the *rpart* package, we defined a classification tree trained on the normalized expression of individual surface markers to partition the cells into subsets with the goal of isolating the cells in the cluster of interest (**Supplementary Information**). We sequentially added gates until including another gate would reduce our sensitivity below 50%. We defined eight populations from every Boolean combination of these three gates: CD26+CD161+CCR6+, CD26-CD161+CCR6+, CD26+CD161-CCR6+, CD26+CD161+CCR6-, CD26-CD161-CCR6+, CD26-CD161+CCR6-, CD26+CD161-CCR6-, CD26-CD161-CCR6-.

### Disease unascertained Boston donor sample processing

PBMCs were isolated from <6hr leukoreduction collars from the Specimen Bank at Brigham and Women’s Hospital using Ficoll-Hypaque as described above. All were discarded samples, collected under IRB protocol 2002P00127, and cryopreserved at 100 million or 50 million cells/mL in 50% FBS, cRPMI, and 5% DMSO in a Freezer buddy.

### Quantification of cytokine production

We thawed aliquots of 150–200 million PBMCs from 3 Boston donors as described above. Samples were washed twice and resuspended in MACS buffer. We magnetically isolated all CD4+ T cells from each sample using a human CD4+ T cell negative isolation kit (Miltenyi biotec, Cat #130-096-533) per manufacturer instructions. We counted CD4+ T cells in each sample after isolation using a Countess™ II Automated Cell Counter and plated each in cRPMI at a concentration of 12.5 million/mL in a 96-well round bottom plate. Cells were incubated overnight at 37°C. After resting the cells, we combined all wells per donor and aliquoted 20 million cells from each donor for flow cytometry staining.

We centrifuged the samples and resuspended each in 1 mL of an antibody master mix consisting of Brilliant Buffer (BD Biosciences) and anti-CCR6 - PECy7 (BioLegend, Clone G03409). We covered in foil and incubated for 25 min at room temperature. Then we added 1 mL of a second antibody master mix consisting of Brilliant Buffer (BD Biosciences) and 9 markers for four-way sorting (**Supplementary Table 12**). We incubated the samples for 25 minutes at 4°C. After staining, we washed the samples twice in 5 mL of MACS buffer, filtered through a 40um mesh filter, resuspended each in 1 mL MACS buffer, and kept all samples on ice in preparation for cell sorting.

We sorted each sample into 4 populations using a BD FACSAria™ Fusion cell sorter. Cells were collected in FACS tubes containing 50% FBS and 50% MACS buffer. Each population was gated on lymphocytes using forward and side scatter and further sorted as follows: 1) Naïve CD4 T cells: CD3+CD4+CD45RO-CD62L+, 2) Other memory CD4 T cells: CD3+CD4+CD45RO+CCR6-/+CD26-/+CD161-/+, 3) Tregs: CD3+CD4+CD25+CD127low, 4) Target population: CD3+CD4+CD45RO+CCR6+CD26+CD161+.

After sorting, we resuspended each sample in cRPMI, counted using a Countess™ II Automated Cell Counter, and plated each sample population at 125,000– 250,000 cells per well in a 96-well round bottom plate. We stimulated one well per population per sample with an equal volume of cRPMI containing a 1:1 ratio of washed CD3/CD28 Dynabeads™ (Thermofisher, Cat #11131D) or with cRPMI for non-stimulated controls. All conditions were incubated overnight at 37°C. After incubation, we transferred the supernatant from each well into a new 96-well round bottom plate and froze the plate at -20°C. We followed manufacturer instructions for the LEGENDplex™ Human Th Panel (13-plex) kit (BioLegend, Cat #740722) in a 96-well V bottom plate. We tested for IL-2, IL-4, IL-5, IL-6, IL-9, IL-10, IL-13, IL-17A, IL-17F, IL-21, IL-22, IFNγ, and TNF. After thawing the supernatants, we diluted our samples 1:10 using Assay Buffer and collected data on a BD LSRFortessa™. We analyzed data using LEGENDplex™ Data Analysis Software. To estimate cytokine concentration in each sorted population, we averaged measurements across 2 technical replicates for each of 3 donors. We compared estimated cytokine concentration between populations with a two-sided t test.

### Intracellular flow cytometry staining

In two experiments, we thawed aliquots of 50–200 million PBMCs from 5 Boston donors and magnetically isolated CD4+ T cells as described previously. We counted the remaining CD4+ T cells in each sample using Trypan blue and a hemocytometer. Then we plated each in cRPMI at a concentration of 10–12.5 million/mL in a 96-well round bottom plate and incubated samples overnight at 37°C. After resting the cells, we recombined all wells per donor and aliquoted 800,000 - 1,000,000 cells into new wells for stimulation. We stimulated cells with an equal volume of 2X PIM (81 nM PMA, 1.34uM ionomycin, and 5 ug/mL brefeldin), and kept the remaining wells unstimulated by adding only an equal volume of 2X brefeldin (5 ug/mL, BioLegend). We incubated plates at 37°C for 4 hours.

After stimulation, we transferred the samples to FACS tubes and washed twice with 500 uL cRPMI. We resuspended the samples in 500 uL of blue fluorescent Live/Dead fixable dead cell stain with PBS (1:1000) (Invitrogen) for 20 min at room temperature. After washing, we resuspended in 50 uL of the first antibody master mix consisting of Brilliant Stain Buffer (BD Biosciences) and CCR6 - PE/Cy7 (BioLegend, Clone G03409). We covered each in foil and incubated for 25 min at room temperature. Then we added 50 uL of a second antibody master mix consisting of Brilliant Stain Buffer (BD Biosciences) and 11 surface markers (**Supplementary Table 12)**. We incubated the samples for 25 minutes at 4°C. After staining, we washed the samples once in MACS buffer.

Next, we followed manufacturer instructions to fix and permeabilize the samples using a Cyto-Fast™ Fix/Perm Buffer Set from BioLegend (Cat #426803). We divided the unstimulated cells per donor in half and stained one tube of unstimulated cells and one tube of stimulated cells per donor with an intracellular antibody master mix consisting of anti-IL17A – APC (BioLegend, Clone BL168) and PE conjugated to either anti-IL-4, anti-IFNy, anti-IL2, anti-IL5, anti-IL9, anti-IL10, anti-IL13, anti-IL17F, anti-IL21, anti-IL22, or anti-TNF in the provided wash buffer (**Supplementary Table 12**). Samples were covered in foil and incubated for 20 min at room temperature. Samples were washed twice in 1 mL of wash buffer and run on a BD LSRFortessa™. Data was analyzed using FlowJo version 10.6.2. The gating structure is shown in **Supplementary Information**.

In two subsequent experiments, we repeated the intracellular staining experiment above using 5 million PBMCs from 8 matched pairs of cases and controls from our Peruvian cohort. We stimulated 500,000 cells per condition and, after extracellular staining, we stained all samples with anti-IL17A – APC (BioLegend, Clone BL168) and anti-IL22 – PE (BioLegend, Clone 2G12A41). We collected data using two BD LSRFortessa™ analyzers.

To compare cytokine production across CD4+ T cell subsets, we gated all CD4+ T cells, all memory CD4+ T cells, all naïve CD4+ T cells, Tregs, and eight populations on Boolean combinations of three surface markers: CD26+CD161+CCR6+, CD26-CD161+CCR6+, CD26+CD161-CCR6+, CD26+CD161+CCR6-, CD26-CD161-CCR6+, CD26-CD161+CCR6-, CD26+CD161-CCR6-, CD26-CD161-CCR6-. We calculated the OR of cells in a given population producing each cytokine, across all donors, with the Cochran-Mantel-Haenszel method. We compared the percent of cells producing IL-17A or IL-22 between cases and controls with a one-sided Wilcoxon signed rank test.

### Data availability

Upon acceptance, all single-cell sequencing data will be made available on GEO. Genotype data will be available on dbGAP.

### Code availability

Upon acceptance, scripts to reproduce analyses will be made available on GitHub.

## Acknowledgments

This work is supported in part by funding from the National Institutes of Health (U19AI111224, UH2AR067677, T32 HG002295, and U01 HG009379, AI049313).

## Author contributions

S.R., D.B.M., and M.B.M. conceptualized and designed the study. A.N. and S.R. designed the statistical and computational strategy and analyzed the data. K.I., S.A., and Y.L. conducted additional statistical analyses. J.I.B., Y.B., S.S., A.N., I.v.R., D.B.M., and S.R. designed the immunoprofiling strategy. J.I.B., Y.B., and S.S. conducted all immunoprofiling experiments. J.J., L.L., and M.B.M. recruited, clinically phenotyped, and obtained blood samples from human subjects. I.v.R., K.L.T. and R.C. organized processing, transportation, and management of PBMCs. C.C., Z.Z., and M.B.M. curated and analyzed clinical phenotype data. A.N. and S.R. wrote the initial manuscript. All authors contributed to the writing and editing the final manuscript.

## Supplementary information

is available for this paper.

