## Extended Data and Supplementary Figures for "Multimodal memory T cell profiling identifies a reduction in a polyfunctional Th17 state associated with tuberculosis progression"

**Supplementary Materials**

Extended Data Figures 1–10

Supplementary Figures 1–9

Supplementary Tables 1–12 (*see .xlsx file*)

### Extended Data

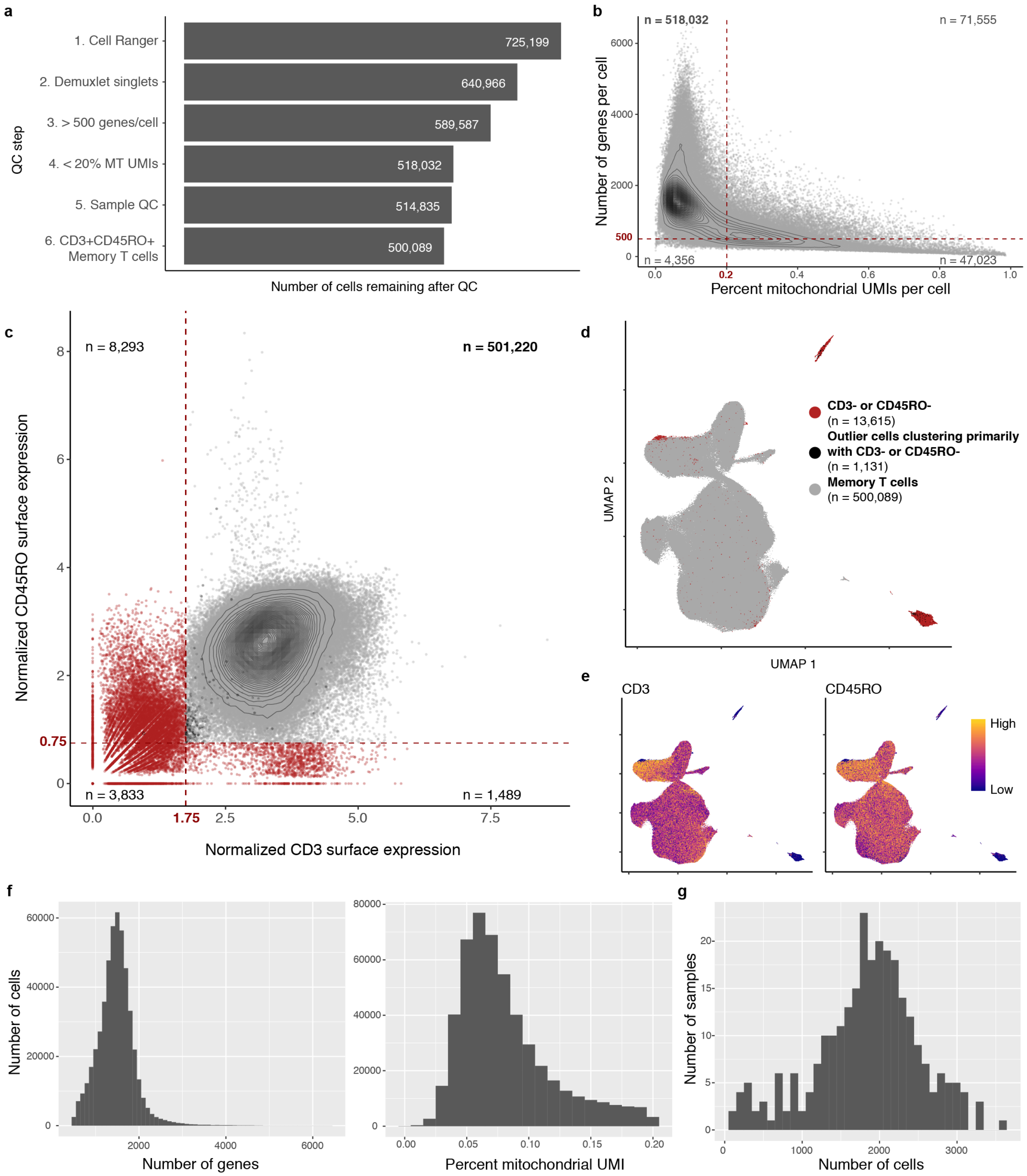

**Extended Data Fig. 1. Quality control of single-cell data. a,** Cell counts over six quality control steps. **b,** Single-cell quality metrics. Each cell is plotted according to the percent of MT UMIs and the number of genes expressed. QC thresholds are demarcated with red dashed lines. Counts indicate the number of cells in each quadrant. **c,** *In silico* memory T cell gating. Each cell is plotted based on its normalized surface expression of CD3 and CD45RO, measured through CITE-seq. Gates are demarcated with red dashed lines. Red cells were removed. Counts represent the number of cells in each quadrant. **d,** UMAP representation of gated cells. Red cells were gated out in (c) and cells clustering with them in the UMAP (shown in black in (c) and (d)) were also removed. **e,** Normalized CD3 and CD45RO surface protein expression. **f,** Distribution of single-cell quality metrics after QC for 500,089 cells. **g,** Distribution of post-QC cell yields for 259 samples.

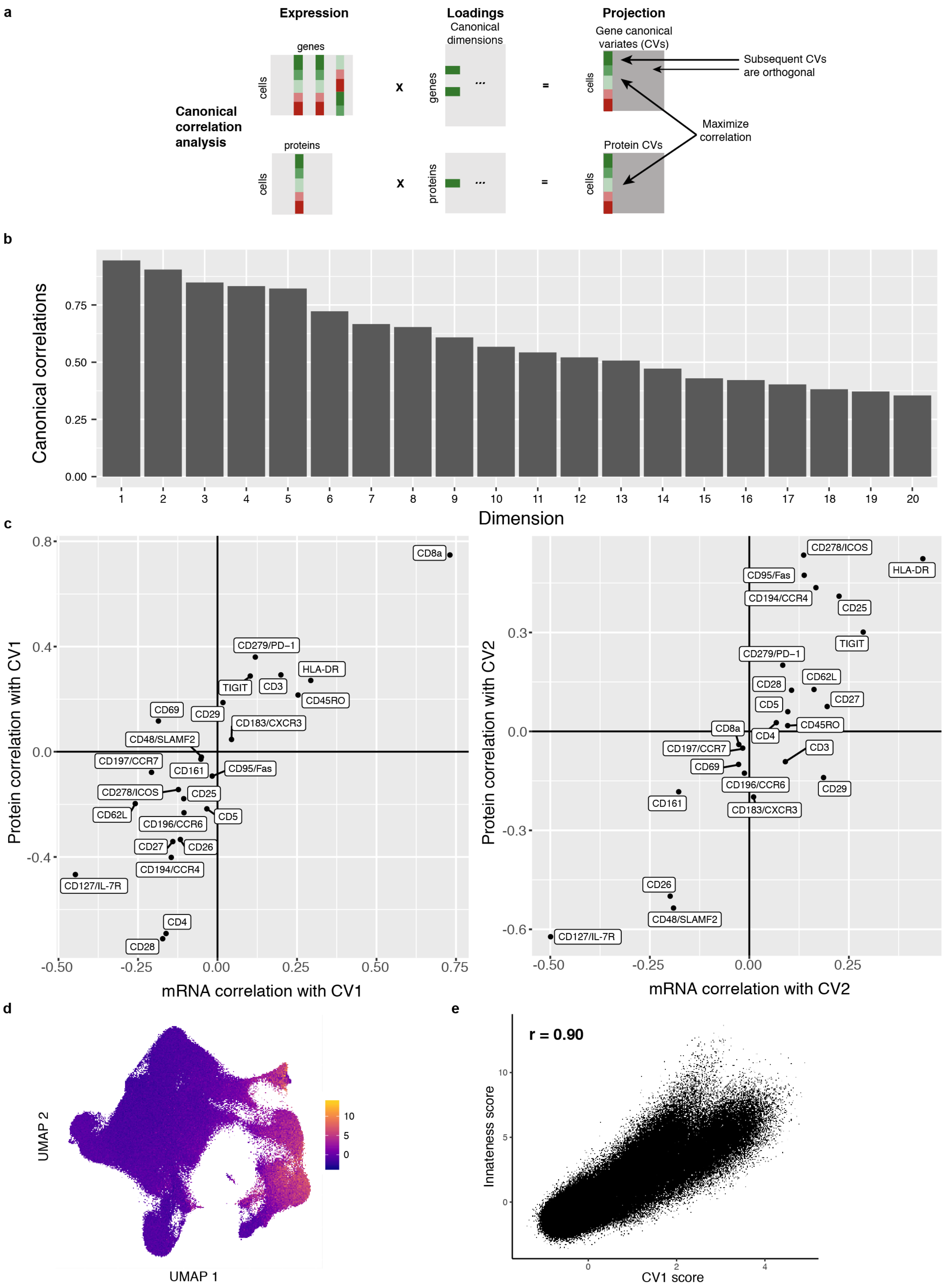
a
Expression
genes
cells
proteins
cells
Canonical correlation analysis
x
x
Loadings
Canonical dimensions
genes
proteins
...
...
=
=
Projection
Gene canonical variates (CVs)
cells
Protein CVs
cells
Subsequent CVs are orthogonal
Maximize correlation
b
Canonical correlations
0.00
0.25
0.50
0.75
Dimension
c
Protein correlation with CV1
mRNA correlation with CV1
0.8
0.4
0.0
-0.4
-0.50
-0.25
0.00
0.25
0.50
0.75
CD8a
CD279/PD−1
HLA-DR
TIGIT
CD3
CD45RO
CD69
CD29
CD183/CXCR3
CD48/SLAMF2
CD197/CCR7
CD161
CD95/Fas
CD278/ICOS
CD25
CD62L
CD5
CD196/CCR6
CD27
CD26
CD194/CCR4
CD127/IL-7R
CD4
CD28
Protein correlation with CV2
mRNA correlation with CV2
0.3
0.0
-0.3
-0.6
-0.50
-0.25
0.00
0.25
CD278/ICOS
HLA-DR
CD95/Fas
CD194/CCR4
CD25
TIGIT
CD279/PD−1
CD62L
CD28
CD27
CD5
CD8a
CD45RO
CD4
CD197/CCR7
CD3
CD69
CD196/CCR6
CD29
CD161
CD183/CXCR3
CD26
CD48/SLAMF2
CD127/IL-7R
d
UMAP 2
UMAP 1
10
5
0
e
r = 0.90
Innateness score
CV1 score
10
5
0
0
2
4

**Extended Data Fig. 2. Multimodal integration with canonical correlation analysis.**
**a,** Schematic of canonical correlation analysis. **b,** Correlations for the top 20 canonical dimensions used in downstream analysis. Bars represent the Pearson correlation between mRNA and protein projections for each dimension. **c,** Marker correlation with canonical variates (CVs). Each marker is plotted based on its mRNA and protein correlation with CV1 (left) or CV2 (right). **d,** “Innateness” scores. UMAP is colored based on a gene expression-derived cytotoxicity score defined in Gutierrez-Arcelus, et al. **e,** Correlation between “innateness” score and CV1. Each cell is plotted based on its “innateness” score from (c) and its CV1 projection, and we report the Pearson correlation coefficient (r).

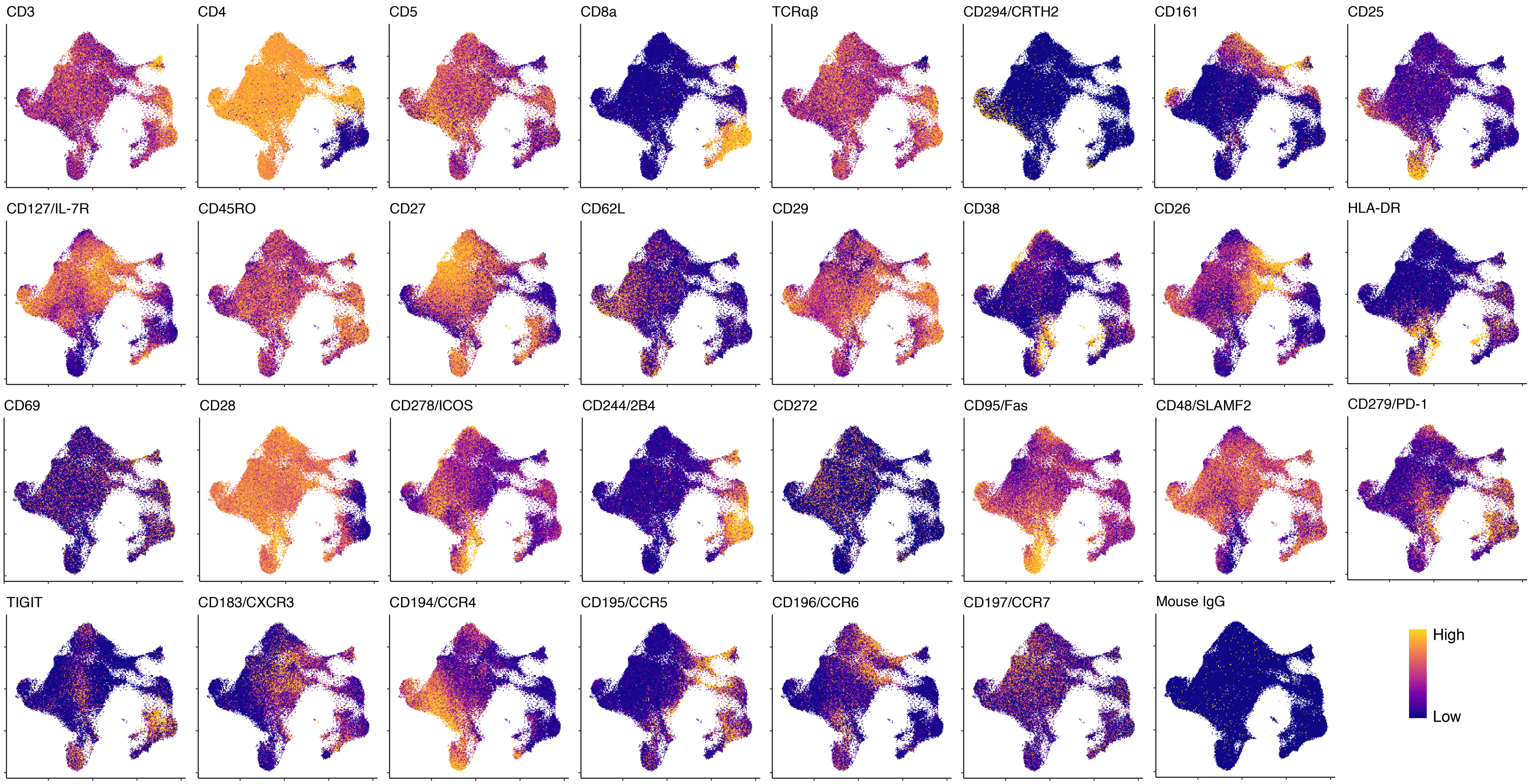

**Extended Data Fig. 3. Single-cell expression of surface markers, measured with CITE-seq.** Colors are scaled independently for each marker from minimum (blue) to maximum (yellow) expression.

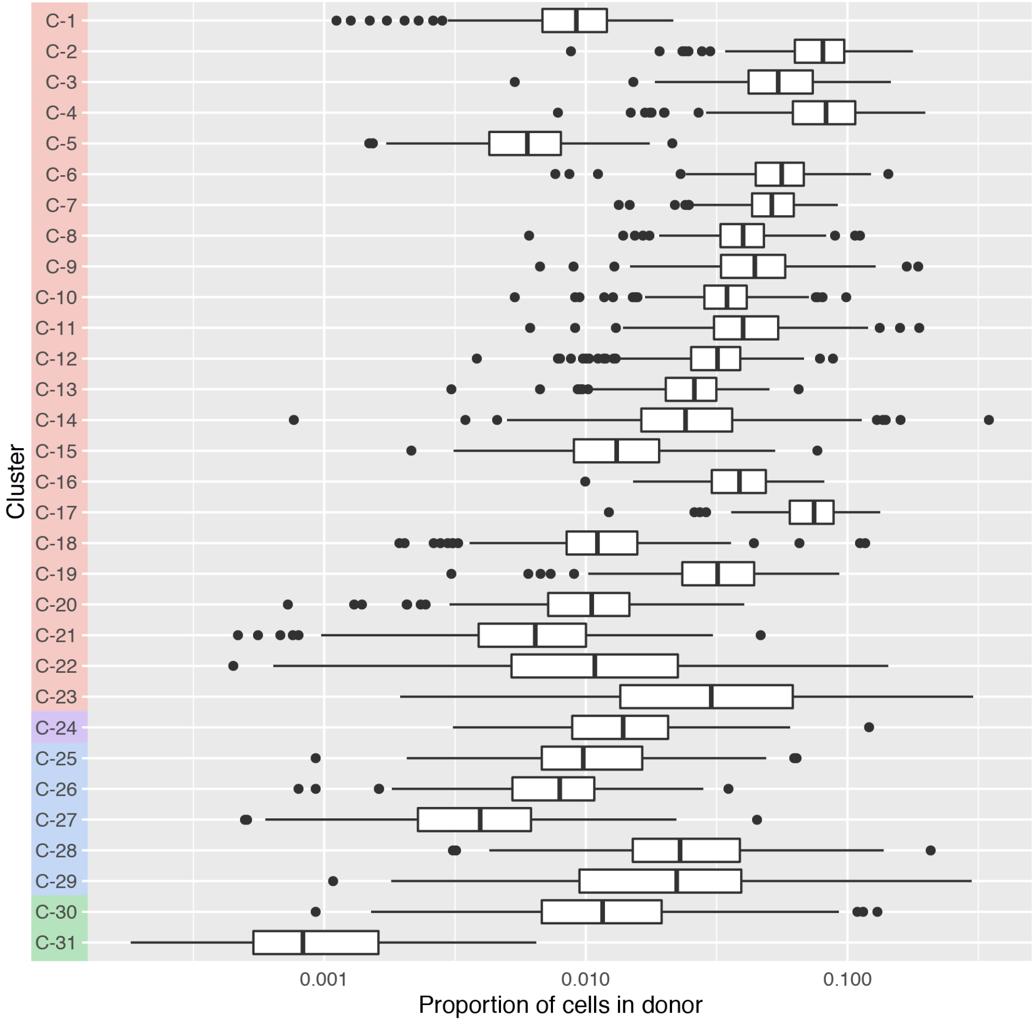

**Extended Data Fig. 4. Cluster proportions across donors.** Boxplots show the distribution of the per-donor proportion of cells in each cluster for 259 donors. Only non-zero proportions are plotted.

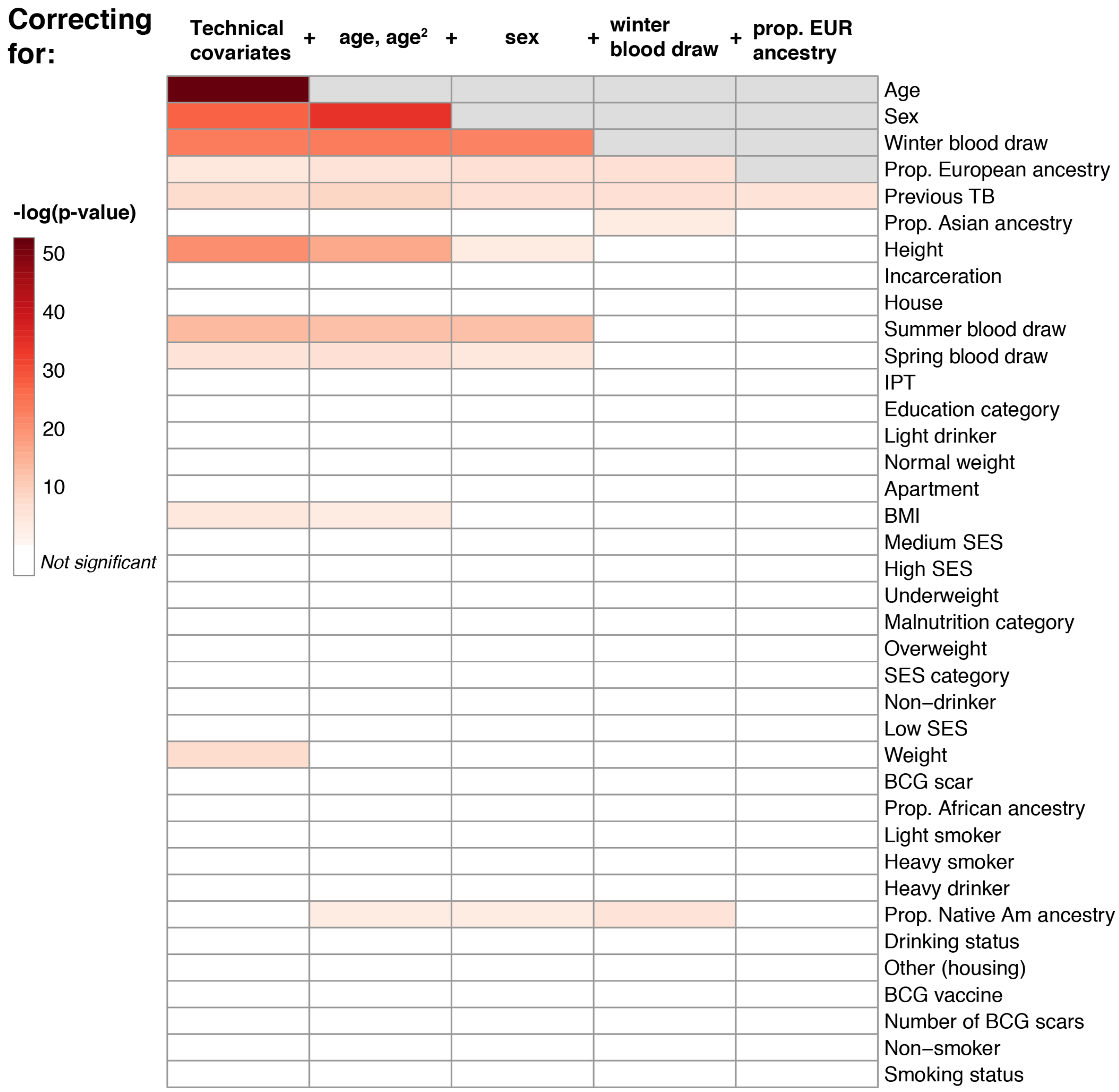

**Extended Data Fig. 5. Associations of covariates with T cell composition.** Each column represents associations from a MASC model fit with the indicated covariate (row) as the contrast, and correcting for the indicated covariates (cumulative column headings, from left) as fixed effects and donor and batch as random effects. Heatmap colors correspond to gamma test p-values, white indicates that the covariate is not significant after multiple testing correction (p > .05/38), and gray indicates that the covariate has already been added to the model. Age and age$^2$ are linear and quadratic terms of age at blood draw. Technical effects are # UMIs/cell and % MT UMIs/cell. EUR = European. IPT = isoniazid preventative therapy. BMI = body mass index. SES = socioeconomic status. BCG = Bacillus Calmette-Guérin.

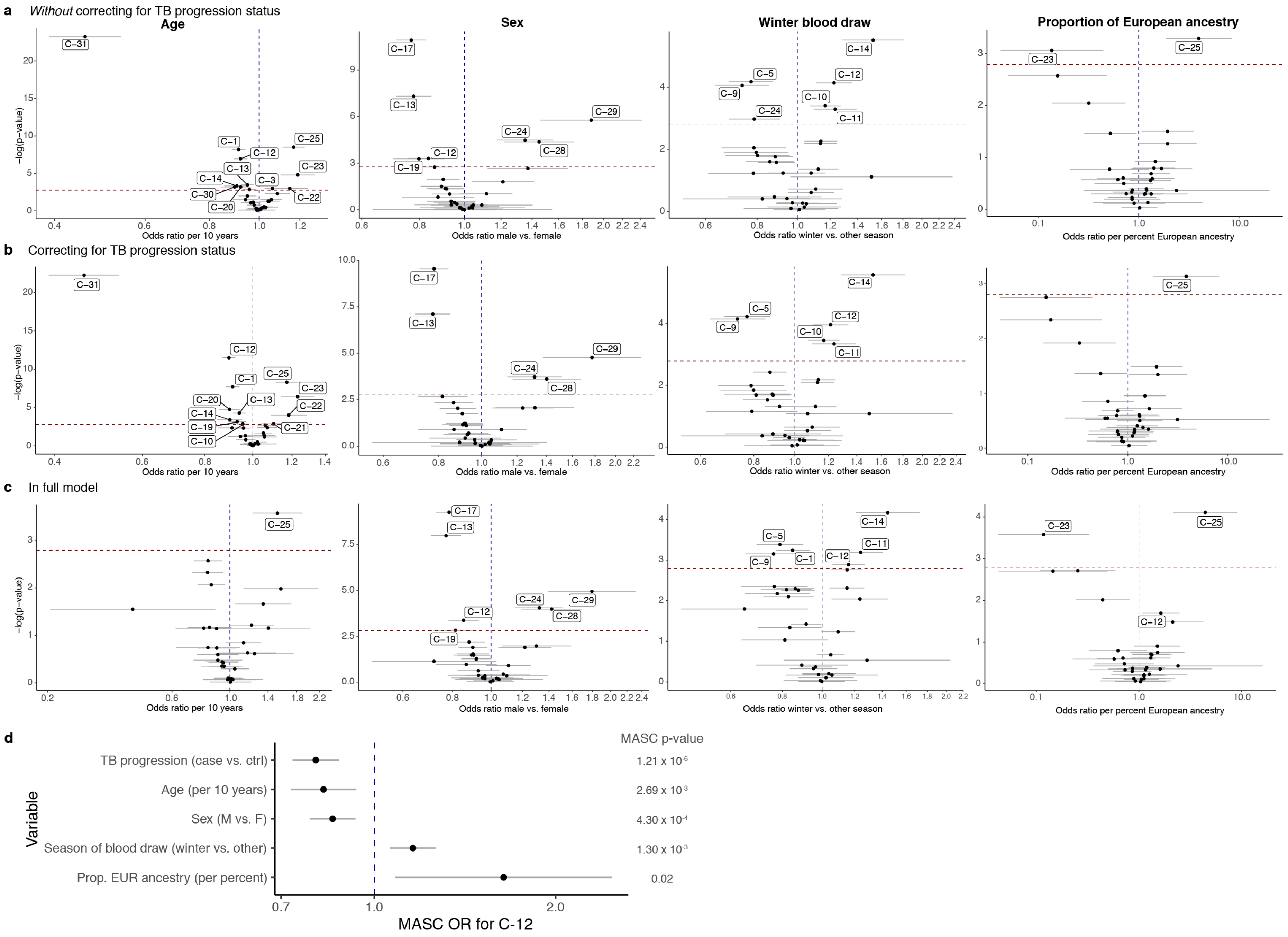
a
Without correcting for TB progression status
Age
Sex
Winter blood draw
Proportion of European ancestry
−log(p−value)
Odds ratio per 10 years
Odds ratio male vs. female
Odds ratio winter vs. other season
Odds ratio per percent European ancestry
b
Correcting for TB progression status
c
In full model
d
TB progression (case vs. ctrl)
Age (per 10 years)
Sex (M vs. F)
Season of blood draw (winter vs. other)
Prop. EUR ancestry (per percent)
Variable
MASC p-value
1.21 x 10-6
2.69 x 10-3
4.30 x 10-4
1.30 x 10-3
0.02
MASC OR for C-12

**Extended Data Fig. 6. Effects of donor covariates on memory T cell states.** Effects of age, sex, winter blood draw, and proportion of European ancestry in **a,** model only correcting for technical covariates (# UMIs/cell, % MT UMIs/cell), donor, and batch, **b,** model in (a) plus TB disease status, and **c,** full model with TB disease status, age, sex, winter blood draw, proportion of European ancestry, technical covariates, donor, and batch. Each cluster is plotted based on the MASC odds ratio of a cell being in that cluster given the contrast covariate, and the -log(LRT p-value) of the association. Error bars show the 95% confidence interval. The red dashed horizontal line corresponds to a Bonferroni-adjusted p-value threshold of 0.05/31 = $1.6 \times 10^{-3}$. Labeled clusters are significant at this Bonferroni-adjusted p-value threshold. **d,** C-12's association with each covariate in the full model. Error bars show the 95% confidence interval.

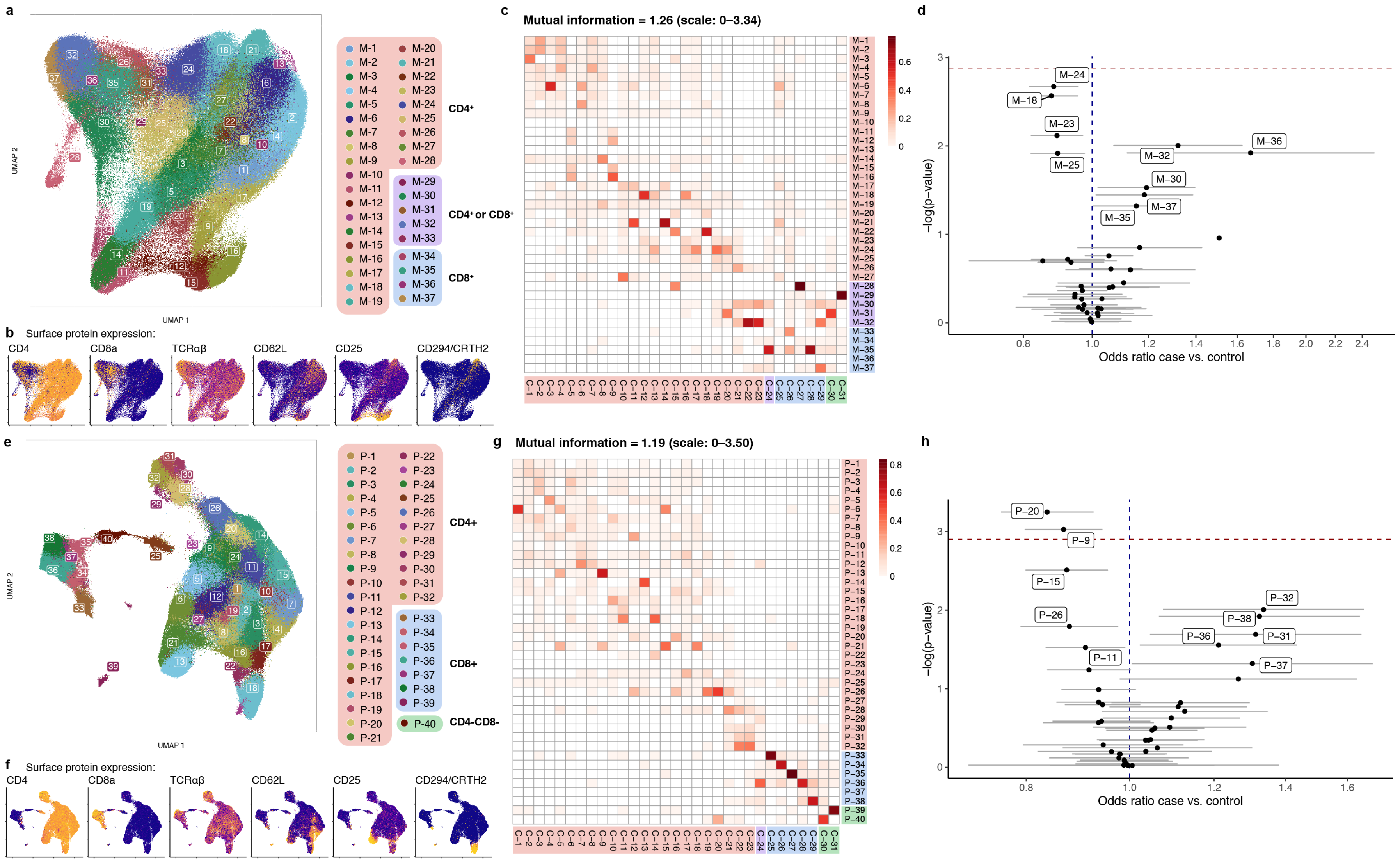
a
b Surface protein expression:
CD4
CD8a
TCRαβ
CD62L
CD25
CD294/CRTH2
CD4⁺
CD4⁺ or CD8⁺
CD8⁺
c
Mutual information = 1.26 (scale: 0–3.34)
d
Odds ratio case vs. control
−log(p−value)
e
CD4+
CD8+
CD4-CD8-
f Surface protein expression:
g
Mutual information = 1.19 (scale: 0–3.50)
h
Odds ratio case vs. control
−log(p−value)
UMAP 1
UMAP 2

**Extended Data Fig. 7. Unimodal clusters and associations with TB disease progression. a-d,** mRNA clusters. **e-h,** protein clusters. **a and e,** UMAPs colored by unimodal clusters. Clusters boxed in red are CD4+, purple are mixed CD4+ and CD8+, blue are CD8+, and green are CD4-CD8-. **b and f**, Expression of major lineage-defining surface markers measured through CITE-seq. The UMAPs are colored by the expression of five markers measured through CITE-seq. Colors are scaled independently for each marker from minimum (blue) to maximum (yellow) expression. **c and g**, Heatmap of overlap between mRNA and multimodal clusters. Colors indicate the proportion of the multimodal cluster (column) overlapping with the mRNA cluster (row). **d and h**, Associations between TB disease status and unimodal clusters. Each cluster is plotted based on the MASC odds ratio of a cell being in that cluster for cases vs. controls, and the -log(LRT p-value) of the association. Error bars show the 95% confidence interval. The red dashed horizontal line corresponds to a Bonferroni-adjusted p-value threshold of $0.05/37 = 1.4 \times 10^{-3}$ for mRNA and $0.05/40 = 1.3 \times 10^{-3}$ for protein. Labeled clusters are significant at a nominal p-value threshold of $p < 0.05$.

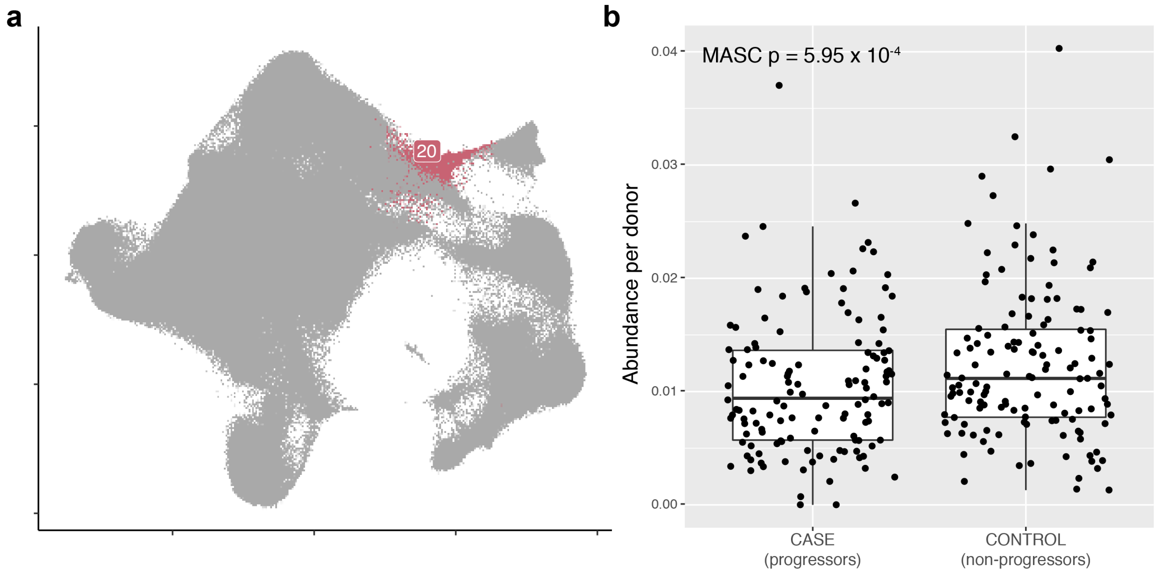

**Extended Data Fig. 8. C-20 is weakly depleted in cases. a,** C-20 highlighted on the UMAP. b, Abundance of C-20 in 128 cases and 131 controls. P-value is from an LRT with 1 d.f. Boxplot center line = median, box limits = 25th and 75th percentile, whiskers = 1.5x interquartile range.

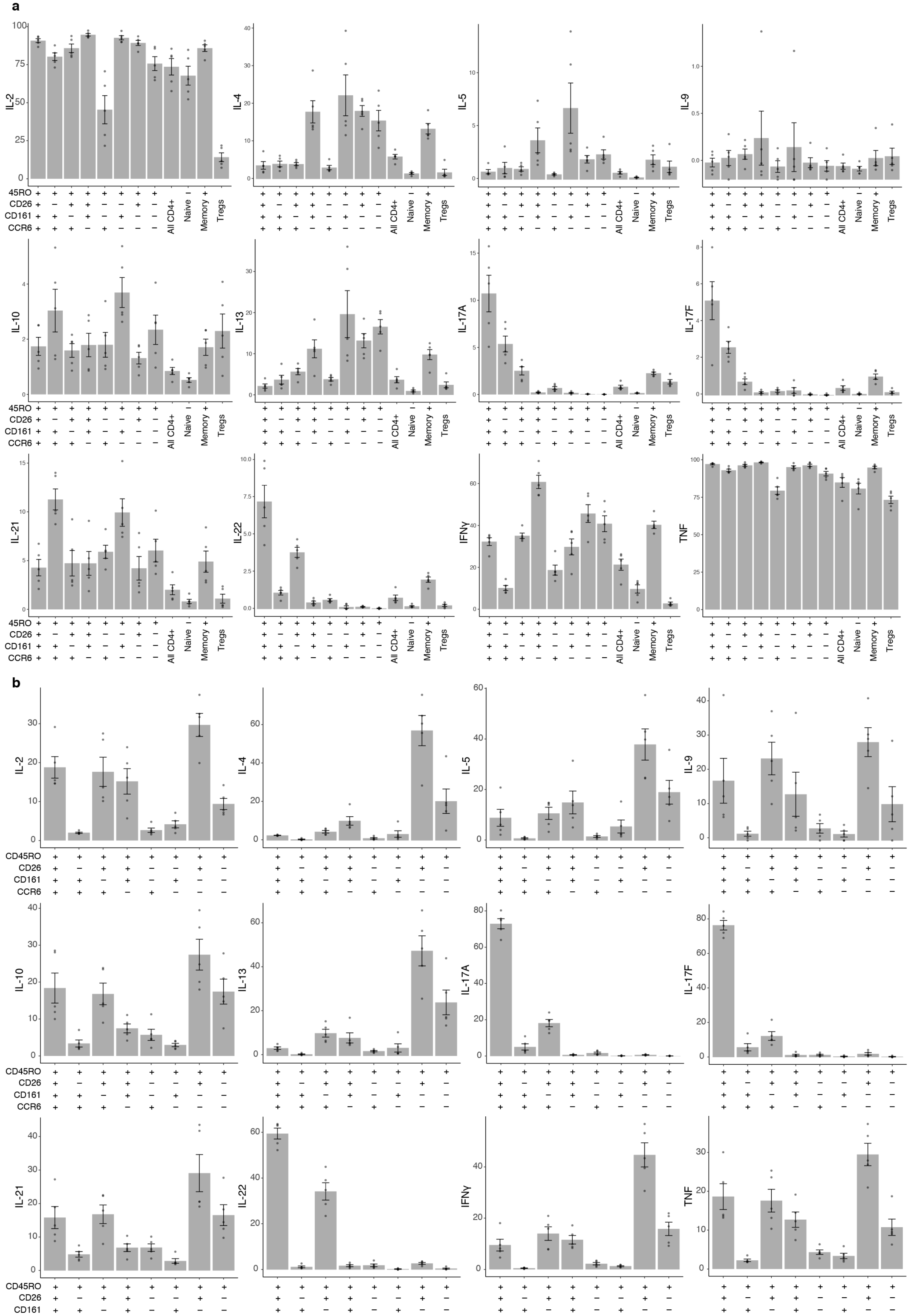
a
IL-2
IL-4
IL-5
IL-9
IL-10
IL-13
IL-17A
IL-17F
IL-21
IL-22
IFNγ
TNF
45RO
CD26
CD161
CCR6
All CD4+
Naive
Memory
Tregs
b
CD45RO
CD26
CD161
CCR6

**Extended Data Fig. 9. Cytokine production in Boston donors.** Bars represent the mean and error bars show standard error of the mean across 5 Boston donors unascertained for TB. **a,** Per-donor percent of cells producing each cytokine in gated populations. **b**, Per-donor percent of total cytokine-producing memory CD4+ T cells in each gated population.

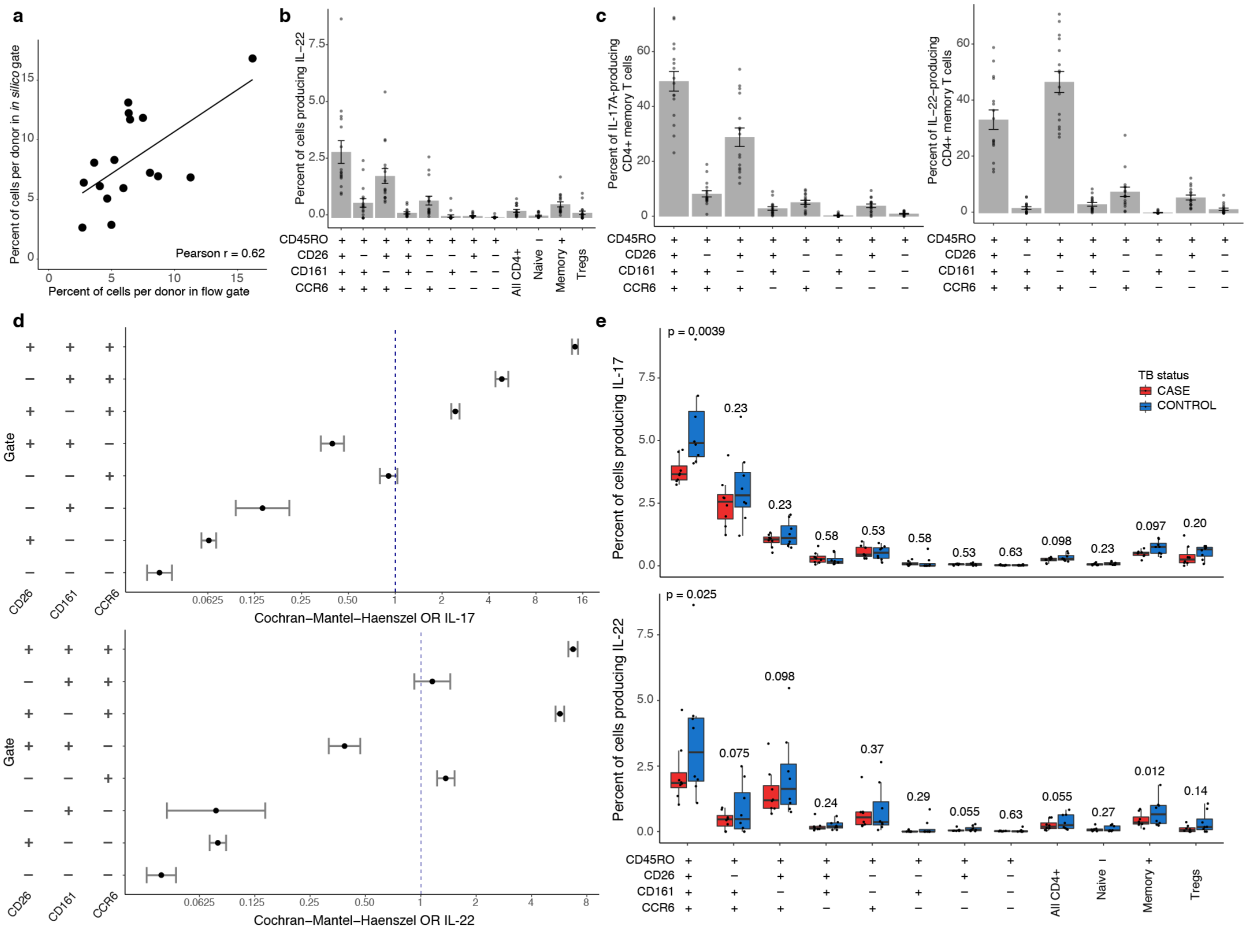

**Extended Data Fig. 10. Cytokine production in Peruvian donors with history of TB infection. a,** Abundance of flow gated and *in silico* gated CD4+CD45RO+CD26+CD161+CCR6+ cells plotted for 16 donors, with a linear best fit line and Pearson correlation coefficient (r). **b,** Per-donor percent of cells producing each IL-22 in gated populations. Bars represent the mean and error bars show standard error of the mean across 16 donors. **c,** Percent of IL-17 (left) or IL-22 (right)-producing cells in gated populations. Bars represent the mean and error bars show standard error of the mean across 16 donors. **d,** Odds ratio of IL-17 or IL-22 production in gated populations. We used the Cochran-Mantel-Haenszel method to calculate the odds ratio of cytokine production inside vs. outside the gated population. Bars show the 95% confidence interval (n = 16). **e,** Case-control comparison of per-donor percent of cells in indicated gates producing IL-17A (top) or IL-22 (bottom). Paired samples (total n = 16) were matched for age, sex, season of blood draw, and proportion of European ancestry. We calculated p-values from a one-sided Wilcoxon signed-rank test. Boxplot center line = median, box limits = 25$^{th}$ and 75$^{th}$ percentile, whiskers = 1.5x interquartile range.

### Supplementary Information

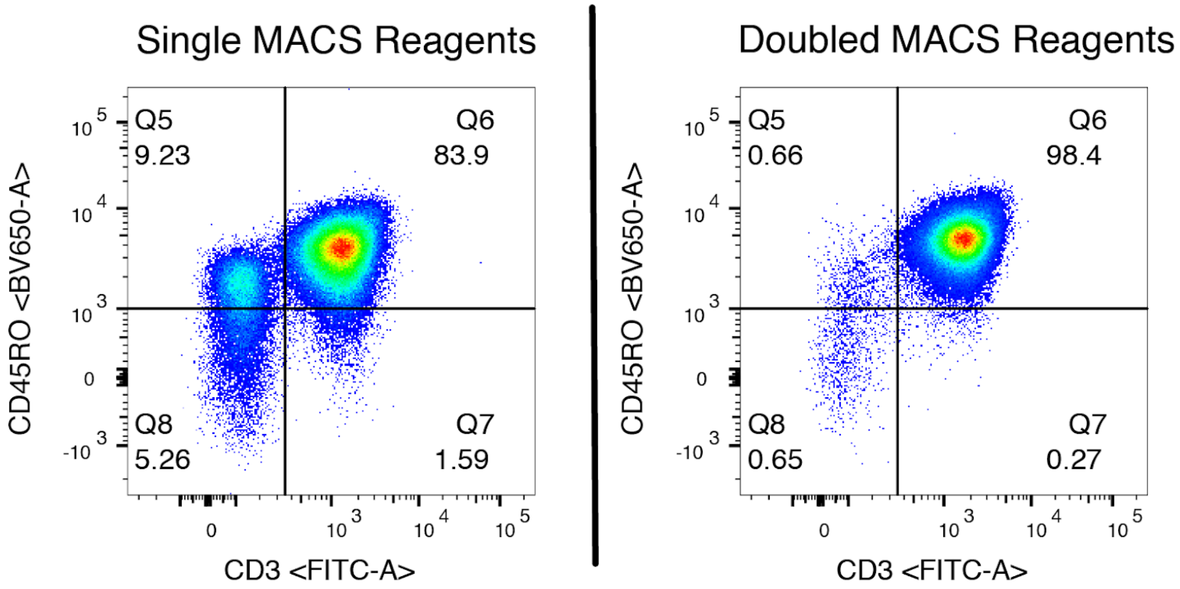

**Supplementary Fig. 1. Memory T cell isolation purity.** Comparison of memory T cell isolation purity with single or doubled MACS reagents. Flow cytometry measurements of CD3 and CD45RO are shown after upstream forward/side scatter, singlet, and live cell gating.

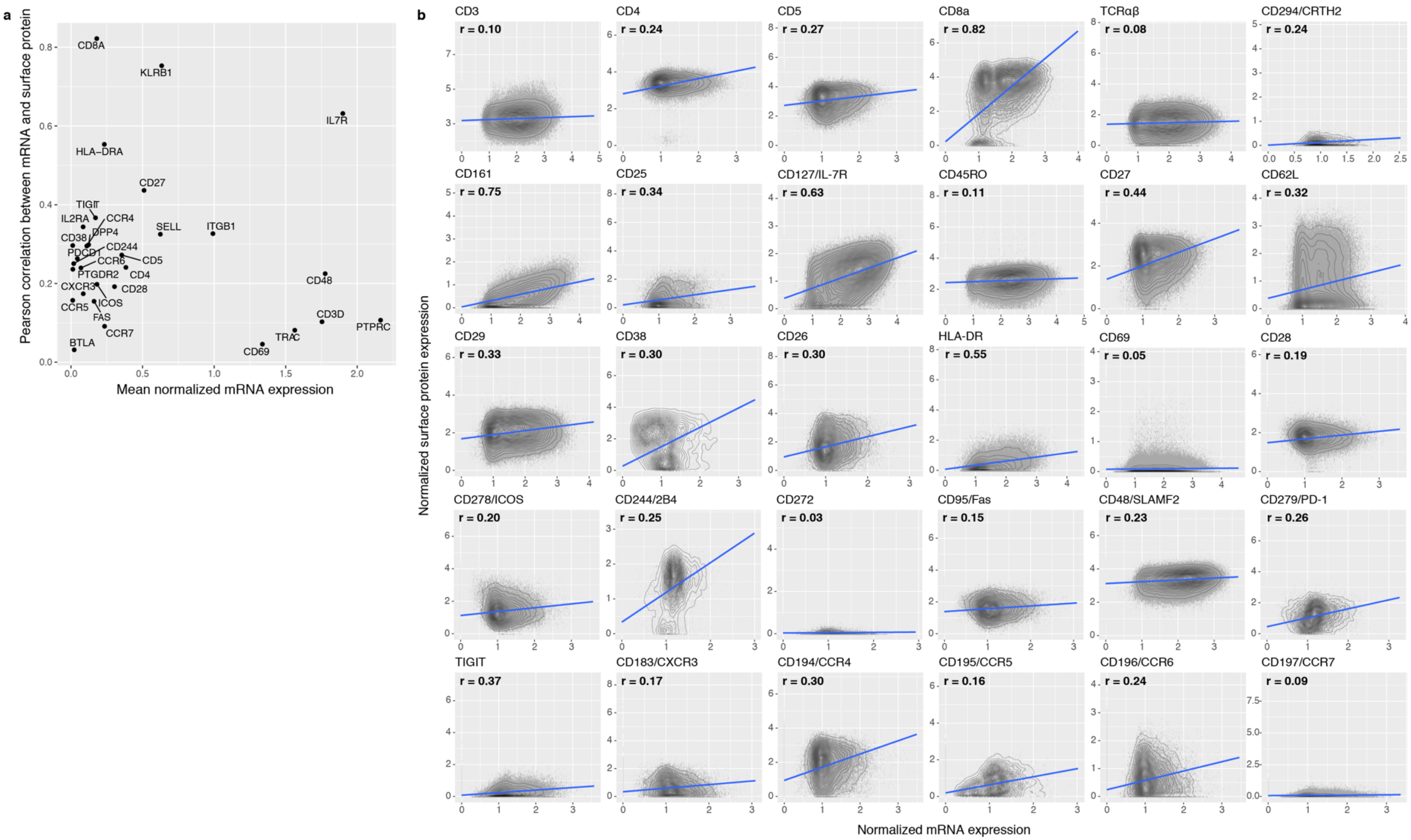

**Supplementary Fig. 2. Correlations between single-cell mRNA and protein expression.** Pearson correlation coefficient (r) was calculated between normalized mRNA and surface protein expression for each marker across cells passing QC. **a,** Pearson correlation coefficient is plotted against average normalized mRNA expression. **b,** Each cell is plotted based on expression of each marker in surface protein and mRNA, both measured through CITE-seq, with density contours. We fit a best-fit line (in blue) with a linear model.

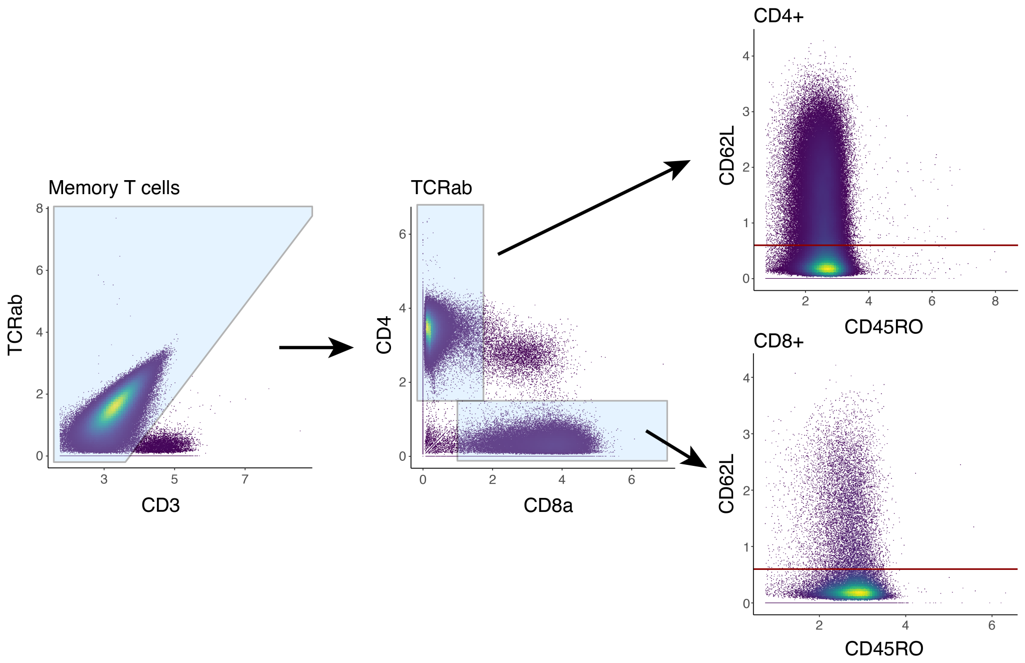

**Supplementary Fig. 3. Representative CITE-seq gating of memory T cell subpopulations.** Biaxial plots show cells passing QC plotted on centered log ratio-normalized protein measurements.

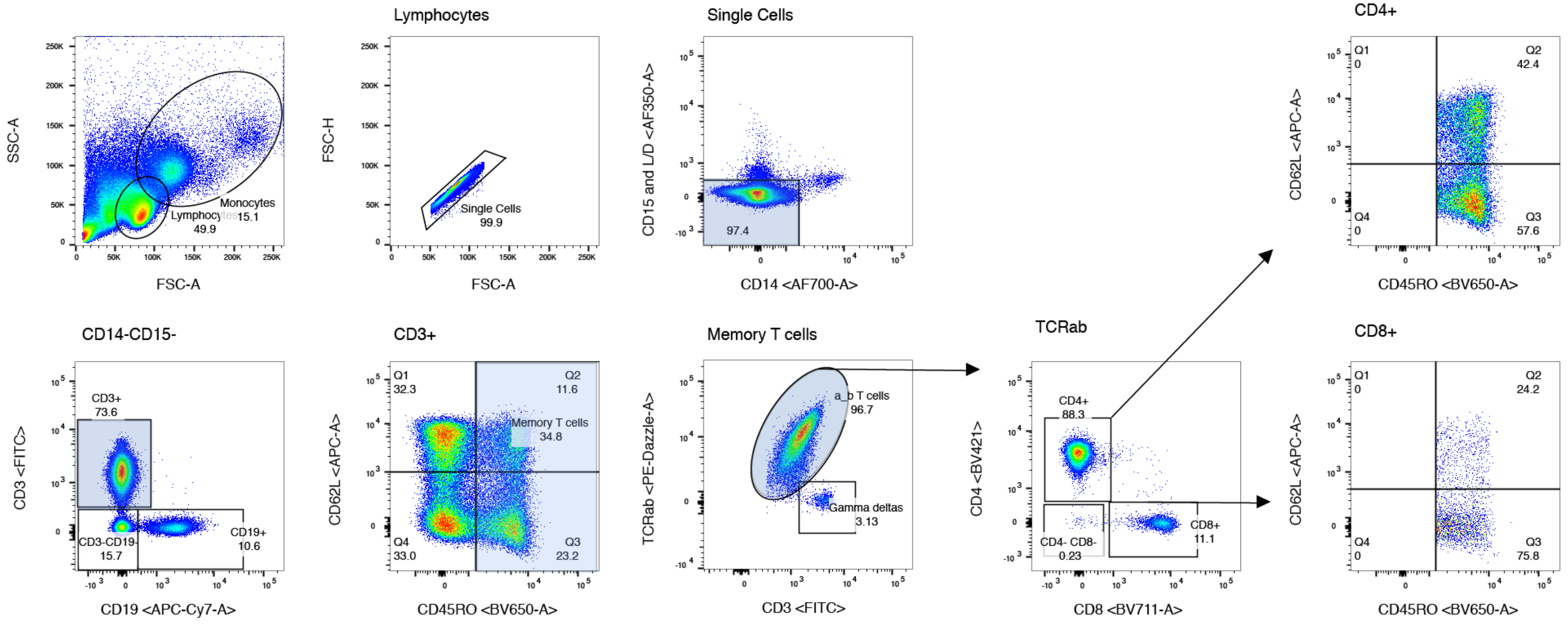

**Supplementary Fig. 4. Representative flow gating of memory T cell subpopulations**

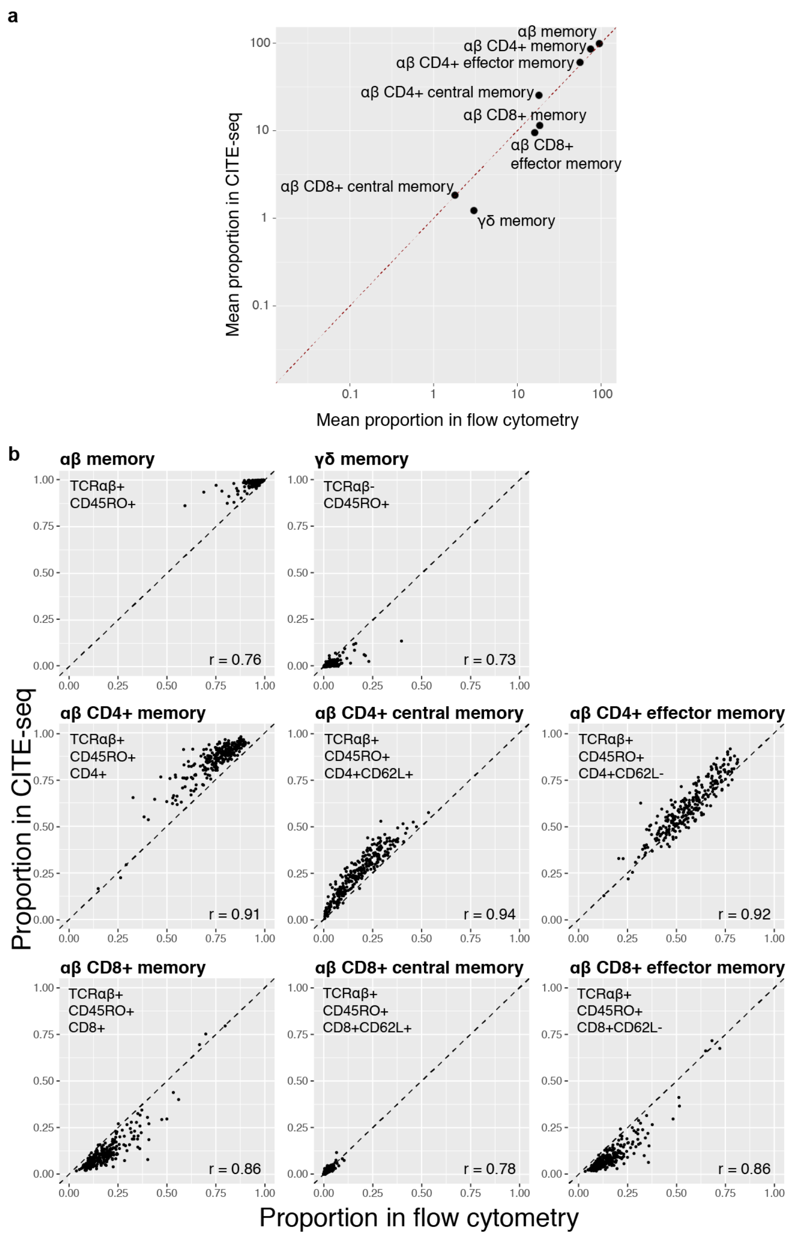

**Supplementary Fig. 5. Comparing proportions of eight major T cell states between flow cytometry and CITE-seq. a,** Average proportions per population in CITE-seq vs. flow cytometry. Gates detailed in (b). Red dashed line indicates a 1:1 relationship. **b,** For each population proportions were plotted across 259 donors. Flow cytometry gating occurred after gating T cells. CITE-seq gating occurred after isolation of memory T cells. The dashed line indicates a 1:1 relationship, and we calculated Pearson correlation coefficients (r) for each state.

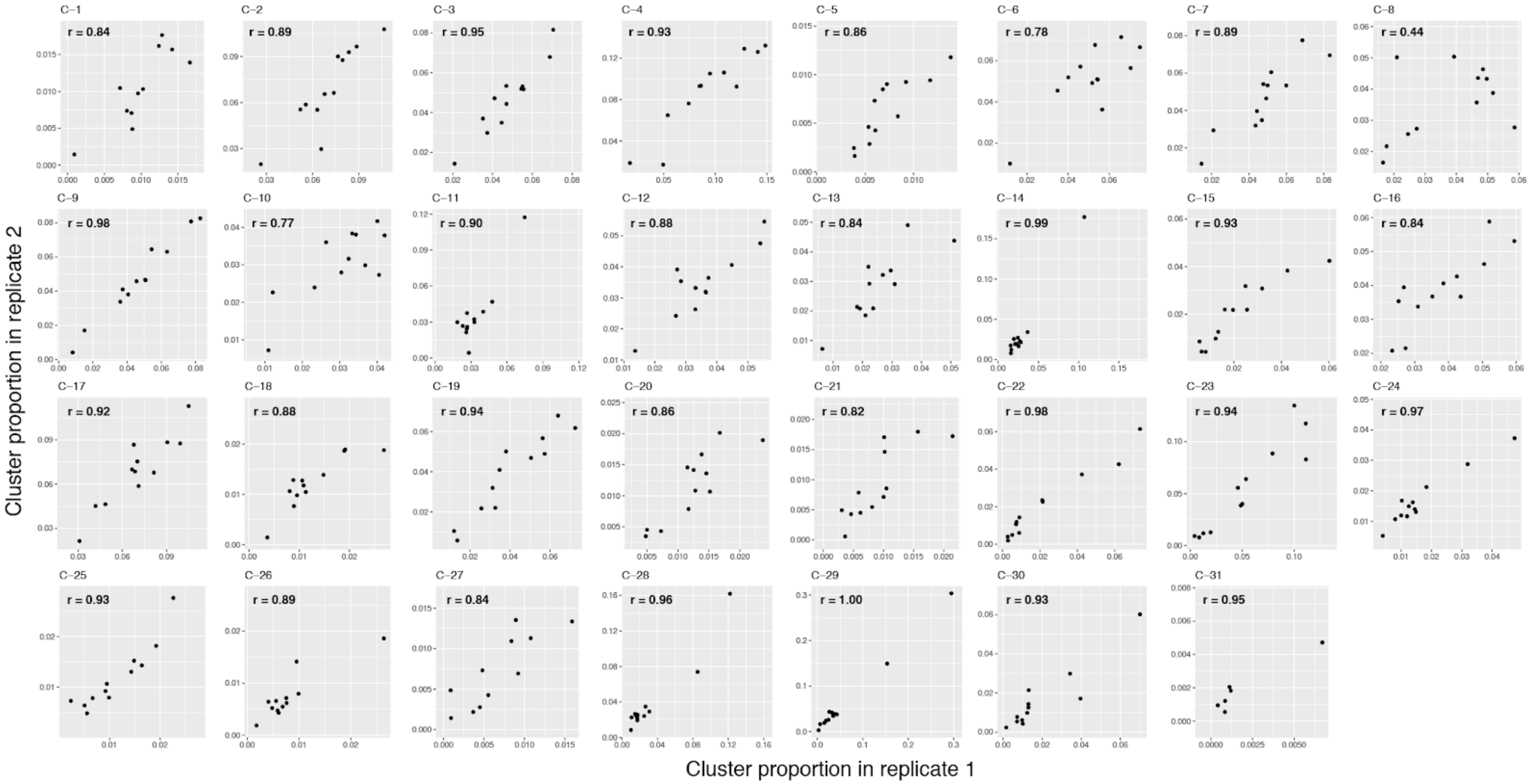

**Supplementary Fig. 6. Technical replicate consistency.** For each of the 31 clusters, we plotted each multimodal donor based on its proportion in replicate 1 and in replicate 2. We calculated the Pearson correlation coefficients (r) for each cluster.

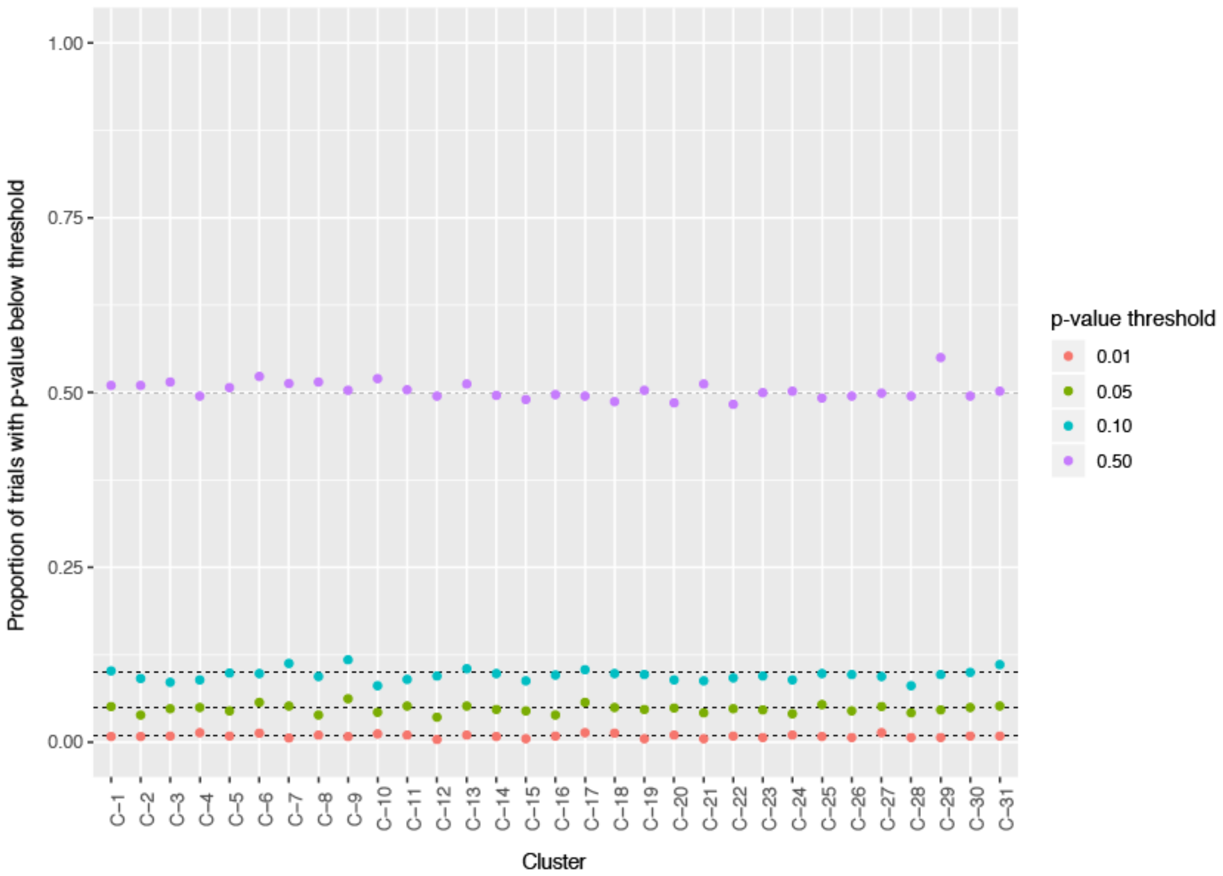

**Supplementary Fig. 7. MASC type 1 error.** We estimated type I error for MASC over 1000 permutations testing cluster associations with permuted TB progression status, adjusting for age, sex, percent MT UMIs/cell (quintiles), and number of UMIs/cell (quintiles). Each model yielded a p-value for each cluster, and in this figure, each point represents the percent of models with a p-value below the indicated threshold for each cluster.

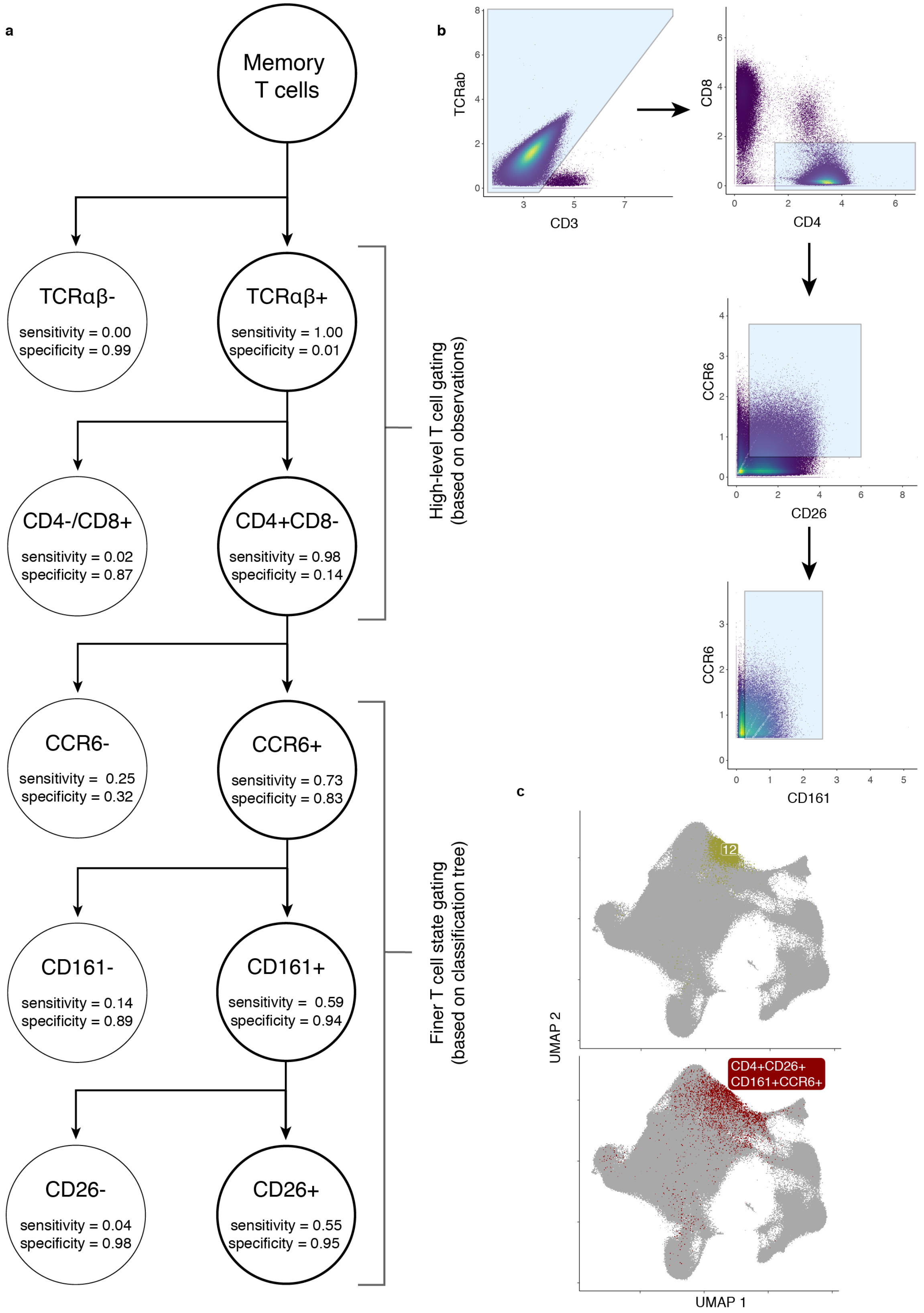
a
Memory T cells
TCRαβ-
sensitivity = 0.00
specificity = 0.99
TCRαβ+
sensitivity = 1.00
specificity = 0.01
CD4-/CD8+
sensitivity = 0.02
specificity = 0.87
CD4+CD8-
sensitivity = 0.98
specificity = 0.14
CCR6-
sensitivity = 0.25
specificity = 0.32
CCR6+
sensitivity = 0.73
specificity = 0.83
CD161-
sensitivity = 0.14
specificity = 0.89
CD161+
sensitivity = 0.59
specificity = 0.94
CD26-
sensitivity = 0.04
specificity = 0.98
CD26+
sensitivity = 0.55
specificity = 0.95
High-level T cell gating (based on observations)
Finer T cell state gating (based on classification tree)
b
TCRab
CD3
CD8
CD4
CCR6
CD26
CCR6
CD161
c
12
CD4+CD26+ CD161+CCR6+
UMAP 2
UMAP 1

**Supplementary Fig. 8. Gating C-12 in CITE-seq data. a,** Classification tree to *in silico* gate C-12. Each level of the tree represents an additional gate. True positives (TP) are cells in C-12 that are in the gate. False positives (FP) are cells not in C-12 that are in the gate. True negatives (TN) are cells not in C-12 that are not in the gate. False negatives are cells in C-12 that are not in the gate. Sensitivity is calculated as TP/(TP + FN). Specificity is calculated as TN/(TN + FP). **b,** Biaxial plots showing gating of TCRab+CD4+CD8-CD26+CD161+CCR6+ cells in CITE-seq data. **c,** Comparison of C-12 and gated population in UMAP space.

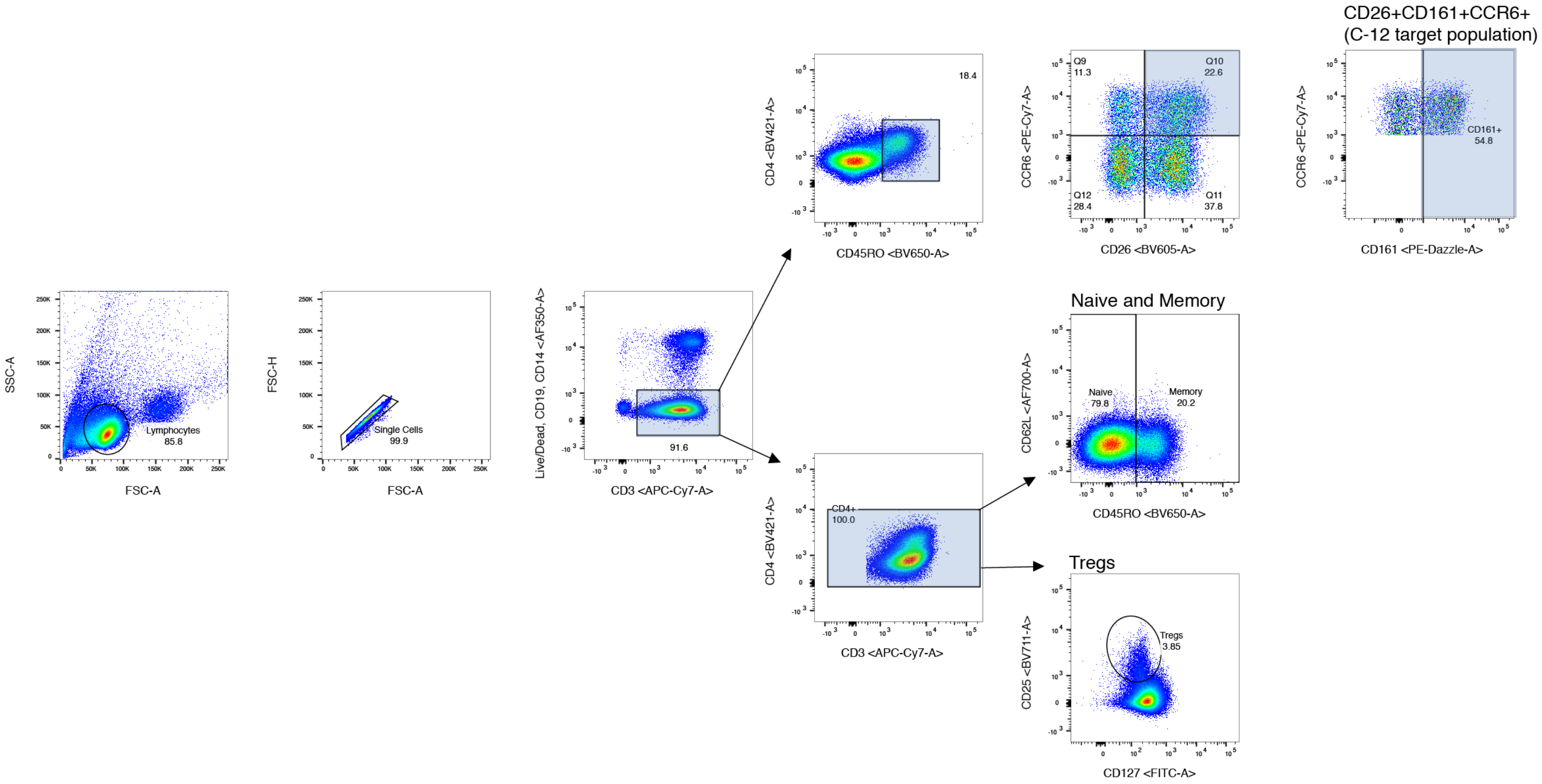

**Supplementary Fig. 9. Representative flow gating of target and control populations from Boston and Peruvian donors.**
